# Discrete photoentrainment of mammalian central clock is regulated by bi-stable dynamic network in the suprachiasmatic nucleus

**DOI:** 10.1101/2024.04.10.588949

**Authors:** Po-Ting Yeh, Kai-Chun Jhan, Ern-Pei Chua, Wun-Ci Chen, Shi-Wei Chu, Shun-Chi Wu, Shih-Kuo Chen

## Abstract

The circadian clock, an evolutionarily conserved mechanism regulating the majority of physiological functions in many organisms, is synchronized with the environmental light-dark cycle through circadian photoentrainment. This process is mediated by light exposure at specific times, leading to a discrete phase shift including phase delays during early subjective night, phase advances during late subjective night, and no shift at midday, known as the dead zone. In mammals, such as mice, the intrinsically photosensitive retinal ganglion cells (ipRGCs) are crucial for conveying light information to the suprachiasmatic nucleus (SCN), the central clock consisting of approximately 20,000 neurons. While the intracellular signaling pathways that modulate clock gene expression post-light exposure are well-studied, the functional neuronal circuits responsible for the three discrete light responses are not well understood. Utilizing *in vivo* two-photon microscopy with gradient-index (GRIN) endoscopes, we have identified seven distinct light responses from SCN neurons. Our findings indicate that light responses from individual SCN neurons are mostly stochastic from trial to trial. However, at the population level, light response composition remains similar across trials, with only minor variations between circadian times, suggesting a dynamic populational coding for light input. Additionally, only a small subset of SCN neurons shows consistent light responses. Furthermore, by utilizing the targeted recombination in active populations (TRAP) system to label neurons that respond to light during early subjective night, we demonstrate that their activation can induce phase delays at any circadian time, effectively breaking the gate that produce photoentrainment dead zone typically observed at midday. Our results suggest the existence of at least two separate time-dependent functional circuits within the SCN. We propose a dynamic bi-stable network model for circadian photoentrainment in the mammalian central clock, where a shifting clock is driven by a dynamic functional circuit utilizing population coding to integrate information flow similar to proposed cortical computational network, rather than a simplistic, consistent linear circuit.

## Introduction

The circadian clock controls many physiological functions in animals, including gene expression, stem cell proliferation, metabolism, neuronal activity, gut microbiota, mood, learning, and memory^1^. In mammals, the suprachiasmatic nucleus (SCN) in the hypothalamus serves as the master clock, orchestrating many peripheral clocks within different tissues ^2–5^. It is essential for these organisms to have an endogenous circadian clock that can be entrained by external environmental cues. The SCN receives direct inputs from a unique group of retinal neurons, the intrinsically photosensitive retinal ganglion cells (ipRGCs), via the retinohypothalamic tract (RHT), conveying environmental luminance information ^6–17^. Consequently, the light pulse signal from ipRGCs is crucial for mammals, such as mice and humans, to synchronize their internal clocks with the external light-day cycle, a process known as circadian photoentrainment^17–21^.

Different species all demonstrate discrete light responses for circadian photoentrainment. Specifically, light exposure during the early subjective night causes a phase delay, while exposure during the late subjective night results in a phase advance ^22–24^. Moreover, during the ’dead zone’ in the middle of the subjective day, light produces only minimal phase shifts ^25^. Recent studies across various model organisms indicate that circadian photoentrainment is regulated by diverse mechanisms. In insects, light directly reaches clock neurons in the brain, allowing cryptochrome or rhodopsin within these neurons to modulate the period-timeless complex and thereby reset the transcription-translation feedback loop (TTFL) in response to external light cues ^26–28^. In contrast, in mammals, ipRGC input to the SCN is required for photoentrainment ^18,20,21^. Light can modulate TTFL components, such as Per1, through glutamate and pituitary adenylate cyclase activating polypeptide (PACAP) released by ipRGCs ^29–34^. While light can induce Per1 expression to daytime levels at many circadian times, this alone does fully explain mammalian circadian photoentrainment. For example, during the day time when light induced phase shift is limited, high-intensity light can still modulate activities or calcium responses within the SCN neurons ^35–37^. Additionally, under distinct light dark cycle, the TTFL coupling and epigenetic profiling of SCN neurons can be dynamic ^38,39^. Finally, the light exposure at different times of day could produce distinct clock genes or immediate early gene expression patterns ^30,40^. Although the molecular mechanism of phase shift is extensively studied and some mathematical models have been proposed ^41–44^, the specific intra-SCN neuronal circuit that are involved in circadian photoentrainment remain poorly understood.

The SCN in mammals is a compact nucleus, composed of many cell types expressing different neurotransmitters with complex intra-nucleus network^45^. Recent single-cell transcriptomic analysis from the mouse SCN suggests it can be divided into at least five distinct molecular subtypes ^46,47^, including vasoactive intestinal peptide (VIP) and gastrin releasing peptide (GRP) expressing neurons in the “core” and arginine vasopressin (AVP) expressing neurons in the “shell” regions ^48^. Classic models have suggested a linear information processing pathway within the SCN. For example, SCN core region, including VIP and GRP neurons, shows high amount of cFos expression than shell region after light pulses ^40,49–51^. Exogenous application of VIP or GRP can produce circadian phase shift both *in vivo* and *in vitro* ^52–55^. Furthermore, silencing the activity of VIP neurons could diminish light-induced phase shift ^36^. Together, these results suggest luminance information is received by VIP and GRP neurons. However, other evidence suggests that the SCN comprises complex networks. For instance, optogenetic activation of VIP neurons can induce phase delay at early subjective night but does not produce the full PRC equivalent to the light pulse ^37^. Conversely, activation of cholecystokinin (CCK) neurons within the SCN can only elicit phase advances during the late subjective night ^56^. Thus, a time-gated functional network could be the underlying mechanism for the discrete properties of the PRC. Given the large number of neurons in the mammalian central clock, it is likely that more complex neuronal interactions and computations are involved in the circadian phase response processes. Consequently, population-wide recording of SCN neurons with single-cell resolution *in vivo* is crucial to decipher the functional network and the neuronal mechanisms behind the circadian photoentrainment.

To elucidate the functional circuitry in the single-cell resolution, we utilized *in vivo* two-photon microscopy with gradient-index (GRIN) endoscopes to record individual SCN neuronal activity using the genetically encoded calcium indicator GCaMP7f ^57,58^. Our findings reveal that SCN neurons display time-gated dynamic network properties, with a small subset of neurons showing consistent light responses at zeitgeber time (ZT) 16 and ZT 22, aligning with circadian phase delay and advance, respectively. Moreover, we could modulate neuronal activity using designer receptors exclusively activated by designer drugs (DREADDs) ^59^ specifically in circadian time (CT) 16 or CT 22 light-responsive SCN neurons with the targeted recombination in the active population (TRAP) mouse system ^60,61^. Activation of neurons trapped at CT 16 could induce phase delays throughout the day, disrupting the photoentrainment dead zone, while activation of neurons trapped at CT 22 could induce phase shift similar to light. Our work suggests that information processing in the SCN likely comprises a bi-stable population-wide dynamic network rather than a linear hierarchical circuitry for circadian photoentrainment.

## Material & method

### Animal

All experiments were performed using C57BL/6 mice raised under 12:12 hr light-dark cycle at room temperature (24°C) under the regulation of National Taiwan University IACUC. Mouse lines employed in the experiments included VIP-Cre (Jackson Labs, strain #031628), Ai14 mice (Jackson Labs, strain #007914) for *in vivo* imaging, and Fos^2A-iCreER^ (TRAP2) mice (Jackson Labs, strain #030323) for behavioral tests. Surgeries were performed at the age of 2 to 8 months, while the in vivo calcium images were obtained at least 4 weeks after GRIN endoscope implantation. The behavioral tests were performed between 4 to 8 months old.

### Stereotaxic Surgeries

All surgeries were performed with a stereotaxic platform and Hamilton syringes controlled by a motorized micromanipulator to ensure precise virus injection and lens implantation to the SCN. Mice were anesthetized by isoflurane inhalation (5% vapor for induction and 1-1.5% vapor for maintenance). For *in vivo* imaging, the GRIN endoscopes (0.6 mm in diameter and 7.4 mm in length, GRINTECH NEM-060-50-00-920-S-1.5P) were implanted after 300 nL AAV9-GCaMP7f (Addgene #104488) injection. The injection and implantation trajectories for GRIN endoscope were rolled 6° clockwise on the coronal plane. Injection needle tips and endoscopes were aimed at anterior-posterior (AP) -0.46 mm and medial-lateral (ML) -0.725 mm from the bregma, and DV -5.2 to -5.4 from the surface of the cerebral cortex. Endoscopes were protected by custom-designed head plates and all materials were secured using dental cement and superbond. For TRAP2 mice in the behavioral tests, 150 nL of AAV9-hSyn-DIO-rM3D(Gs)-mCherry or AAV9-hSyn-DIO-EGFP (Addgene #50458 and #50457) were bilaterally injected, aimed to AP -0.46 mm and ML ±0.2 mm from the bregma and DV -5.76 from the surface of the cerebral cortex.

### In vivo Deep Brain Calcium Imaging

A custom-built two-photon microscope with 23 Hz frame rate at 512x512 pixel resolution was employed to acquire calcium images. GCaMP7f and tdTomato were excited using a 920 nm pulse laser (Coherent, US) at 90 mW output measured after a Zeiss 20x/0.5 objective, whose numerical aperture matches that of the GRIN endoscope. Awake mice were mounted on a custom designed two-dimensional tilting stage during the image acquisition sessions. A free-rotating wheel was set for mice to relieve stress from the physical constraints of their heads. To stimulate retinal photosensitive cells, a 488 nm light diffused onto a 10 x 26 mm screen in front of the eyes of mice. Average power of light on the screen was 1.76 μW/mm^2^. GCaMP7f signal were detected using 525/39 nm bandpass filter, and tdTomato signal were detected using 625/90 nm bandpass filter. Every recording session containing 3 trails for each focus plane, and is at least 14 hours apart from each other to avoid excessive photobleaching.

### In vivo Calcium Image Data Processing

To examine the dynamic responses of individual SCN neurons, we followed the outlined workflow ^62^: Firstly, we corrected motion artifacts in the acquired image slices induced by mice movement ^63^. Next, we utilized Cellpose to segment cell regions, creating cell masks that enable the extraction of calcium traces from the acquired sequence of x-y image slices ^64,65^. Furthermore, to eliminate the decrease in fluorescence intensity caused by photobleaching^66,67^, we applied curve fitting to the time series data of each neuron, removing the decay trend ^68,69^. Finally, to make the fluorescent intensity of each neuron from all trials comparable, we standardized the intensity using the normalized Z-score method. Further details on each step are provided as follows.

Motion correction: As per the mechanics of our mouse imaging platform, motion during the imaging process can be regarded as horizontal movement devoid of shearing and rotation. Consequently, motion correction can be attained through translation transformation ^70^:

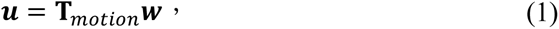

where ***w*** = [*w_x_ w_y_* 1]*^T^* and ***u*** = [*u_x_ u_y_* 1]*^T^* are the space coordinates of a voxel before and after correction, respectively. *T_motion_* is the transformation matrix:

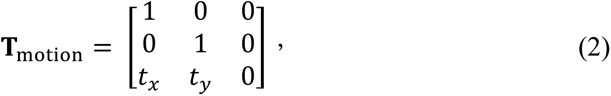

where the parameters *t_x_* and *t_y_* denote the displacement along the x-axis and y-axis of the image slices, respectively. These parameters can be obtained through optimization algorithms in conjunction with a similarity measure *C* (Mattes mutual information metric):

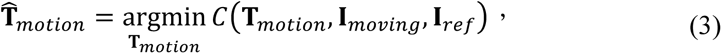

Here, **I***_ref_* represents the reference image, and **I***_moving_* denotes the image requiring correction. The optimization algorithm employs a regular gradient descent method for iterative optimization. The image at the start of the sequence serves as the reference image. After estimating the matrix **T***_motion_* it is used to correct motion in the image **I***_moving_*.

Cell segmentation and extraction of calcium traces: Following the removal of motion-induced artifacts, we calculated the average of all image slices and employed the resulting mean slice as input for cell segmentation, utilizing Cellpose. We use the pre-trained model “CP” provided by Cellpose, with the cell diameter parameter set to 25 pixels. Subsequently, the segmented cell regions were labeled. Considering potential slight variations in activated neurons across different experiments, we manually adjusted their indices to ensure the consistent identification of the same group of neurons. Finally, we computed the average trace of a specific neuron by utilizing the traces of the voxels within the region associated with that neuron.

Detrend: We utilized the trust region optimization method, which is a nonlinear least squares approach, to estimate the decay coefficient for each calcium trace. Subsequently, the decay trend was subtracted to produce a detrended calcium trace, which was then utilized for further analysis. Moreover, background intensity extracted from the out-of-endoscope pixels was also subtracted in this step to exclude those slight (<50 of intensity unit. Soma of neurons were mostly 4000-15000 units in comparison) yet detectable signal changes caused by stimulation light and other residual signal fluctuations in the acquisition system.

Normalization: To perform further analysis, we transformed the fluorescent intensity of each neuron from each trial using a Z-score-based method:

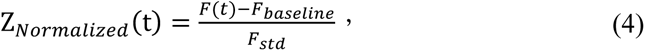

where Z_Normalized_(t) represents the normalized Z-score of assigned time (frame), F(t) represents the fluorescent intensity of assigned time after detrending, F_baseline_ represents the median intensity of the baseline, and F_std_ represents the standard deviation of intensity across the 120-second recording. Instead of using the classic Z-score that defines the average of all values as zero, we set the median of the 30-second baseline as zero to better illustrate and compute the light-evoked neuronal responses.

### Behavioral Phase Shift

To trap light-induced cFos-positive cells, stock solution of 4-hydroxytamoxifen (4-OHT; Sigma, Cat# H6278) was prepared at 100 mg/mL in ethanol and stored at –80°C for up to one month. Before usage, the stock solution of 4-OHT was mixed with sunflower seed oil (Sigma, Cat #s 259853) to final concentration at 10 mg/mL. Immediately after a 10-min 900-lux light pulse at CT 16 or CT 22, TRAP2 mice were intraperitoneally (IP) injected with 4-OHT at 500 mg/kg dosage. After at least 2 days, mice were transferred to wheel-running cages and housed in LD with *ad libitum* food and water. After at least 7 days of free-running in constant darkness (DD), mice were IP injected with CNO at 1 mg/kg or saline at CT 2, CT 6, CT 8, CT 16, and CT 22 for chemogenetic manipulation. Clozapine-N-oxide (CNO, Sigma, Cat# C0832) solution was prepared at 0.2 mg/mL in saline. Phase shifts were determined by using activity onset analysis in Clocklab (Actimetrics, US). Phase shifts were assessed by calculating the time difference observed on a subsequent day after each drug injection to the regression lines. The days of drug administration were excluded from the analysis.

### Immunofluorescence

Mice were anesthetized with tribromoethanol (avertin, 250 mg/Kg) and perfused with 10 ml cold PBS followed by 40 ml 4% PFA for fixation. Brains were isolated and postfixed in 4% PFA for 24 hours. Serial 80 μm coronal brain slices were obtained by vibratomes. Sections were blocked with 10% normal goat serum in PBS containing 0.2% Triton-X and incubated overnight at 4 °C with primary antibodies. Following the wash, sections were incubated for 2 h with fluorescently labeled secondary antibodies. Brain sections were stained with anti-cFos mouse monoclonal antibody (1:1000, Abcam, ab208942), anti-vasopressin rabbit antibody (1:1000, Immunostar, 20069), and anti-GFP chicken antibody (1:1000, Abcam, ab13970). Secondary antibodies used were goat anti-mouse IgG Alexa 488 (Invitrogen, A21121, 1:500), goat anti-rabbit IgG Alexa 633 (Biotium, 20123, 1:500), and goat anti-chicken IgG Alexa 633 (Invitrogen, A21103, 1:500). Sections were then mounted with DAPI-containing Rapiclear (SUNJin lab, Taiwan) and visualized by a Leica confocal microscope (SP5).

## Results

### In vivo Two-Photon Calcium Imaging Reveals Diverse and Dynamic Light Responses in SCN Neurons

Current methodologies, such as *in vivo* photometry and *in vitro* electrophysiology, provide limited insights into how the SCN processes light information for photoentrainment, due to their lack of single-cell resolution or *in vivo* applicability, respectively. To advance our understanding of the SCN’s photoentrainment mechanism, we employed two-photon microscopy with GRIN endoscopes in awake mice, targeting the SCN to observe real-time neuronal activity at the single-cell level using a calcium imaging approach (Figure 1A). VIP-Cre; flex-tdTomato mice were injected with AAV9-hSyn-GCaMP7f into the SCN (Figure 1B and Extended Figure 1A) and subsequently maintained under a 12:12 light-dark cycle for recordings at ZT 8, ZT 16, and ZT 22. This *in vivo* setup allows us to track single neurons across different ZT time points longitudinally (Figure 1C). Furthermore, two-photon system also allow us to image across several hundred microns in dorsoventral axis (Figure 1D) compared to single photon system ^71^. Here, each recording session consisted of six trials of a 30-second dark baseline followed by a 90-second light exposure (488 nm, 1.76 μW/mm2) to the eyes. Activity from two z-positions within the SCN, separated by 105 µm, was captured during the same session (Figure 1E). After motion correction and cell segmentation, we successfully identified 338 regions of interests (ROIs) from three mice across all sessions (Extended Figure 1C-D). Among these ROIs, 113 neurons, including 46 tdTomato-positive VIP neurons (40.7%) were consistently registered across 27 sessions (Figure 1F). Furthermore, of the neurons located in the ventral focus plane, 44.6% are VIP^+^, while in the dorsal focus plane, 33.3% are VIP^+^. Subsequent analysis of the average neuronal response indicated a modest, yet consistent, positive light-induced response among SCN neurons at all three ZTs (Figure 1G), confirming prior photometry findings ^37^.

**Figure 1.**
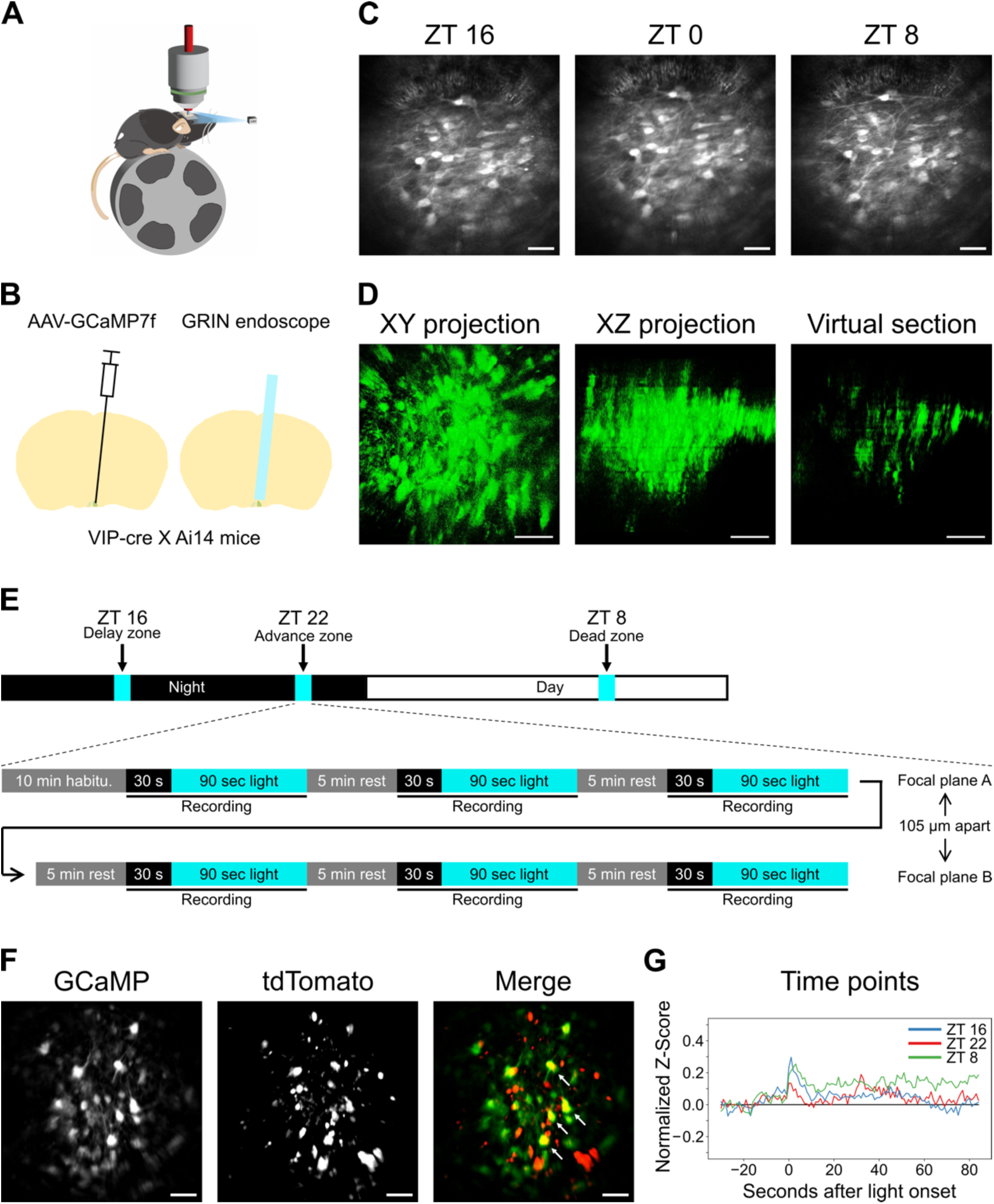
Observation of SCN neuronal light responses in awake mice. **A-B.** An illustration of recording setup and surgical procedure. **C.** Representative two-photon *in vivo* GCaMP images (averaged) of SCN taken at different ZT times through the GRIN endoscope. **D.** Representative 3D-projection of SCN GCaMP signal using z-stack images through the GRIN endoscope. The XY projection shows the summation of horizontal planes, while the XZ projection shows the summation of sagittal planes of the SCN. The XZ virtual 45-μm-thick section demonstrates the elongated point spread function in the z-axis through GRIN endoscopes. **E.** Schemes for time points experiments (upper panel) and calcium imaging recording (lower panel). **F.** Representative images of GCaMP and tdTomato signals collected simultaneously with fast-scanning two-photon system. **G.** Averaged light-response of identified neurons at 3 ZT times. Red: ZT 22, green: ZT 8, blue: ZT 16. n = 3 animals. Scale bars are 100 μm, according to the imaging side of the GRIN endoscope.

To dissect individual SCN neuronal light responses, we analyzed the Z-score of GCaMP signals from 113 neurons (Figure 2A and Extended Figure 2). Given the diversity of spontaneous calcium transients exhibited by SCN neurons, we averaged the Z-scores in 5-second bins to analyze light response trends for 90 sec. Subsequently, principal component analysis (PCA) was conducted on data from all trials of the 113 neurons, and k-means clustering was utilized to categorize the light responses into seven distinct clusters of light response (Figure 2B). We plotted the mean light response for each cluster, including transient activation (cluster 1, 7.48%), sustained activation (cluster 2, 5.01%), delayed activation (cluster 3, 14.7%), marginal response (cluster 4, 35.5%), delayed inhibition (cluster 5, 16.1%), transient inhibition (cluster 6, 16.8%), and sustained inhibition (cluster 7, 4.42%) (Figure 2C). Notably, neurons were not limited to a single type of light response; on average, each neuron exhibited 5.57 ± 1.08 (mean with SD) distinct types of light response across the 27 trials (Figure 2D and 2E). Furthermore, even when using the most stringent criteria that consider marginal responses (cluster 4) as non-responsive to light, every neuron exhibited either activation (cluster 1-3) or inhibition (cluster 5-7) in at least 4 out of the total 27 trials (16%). These findings not only demonstrate the capability of SCN neurons to generate multiple types of light response but also imply the presence of a dynamic functional circuitry involving most if not all SCN neurons to encode light signals.

**Figure 2.**
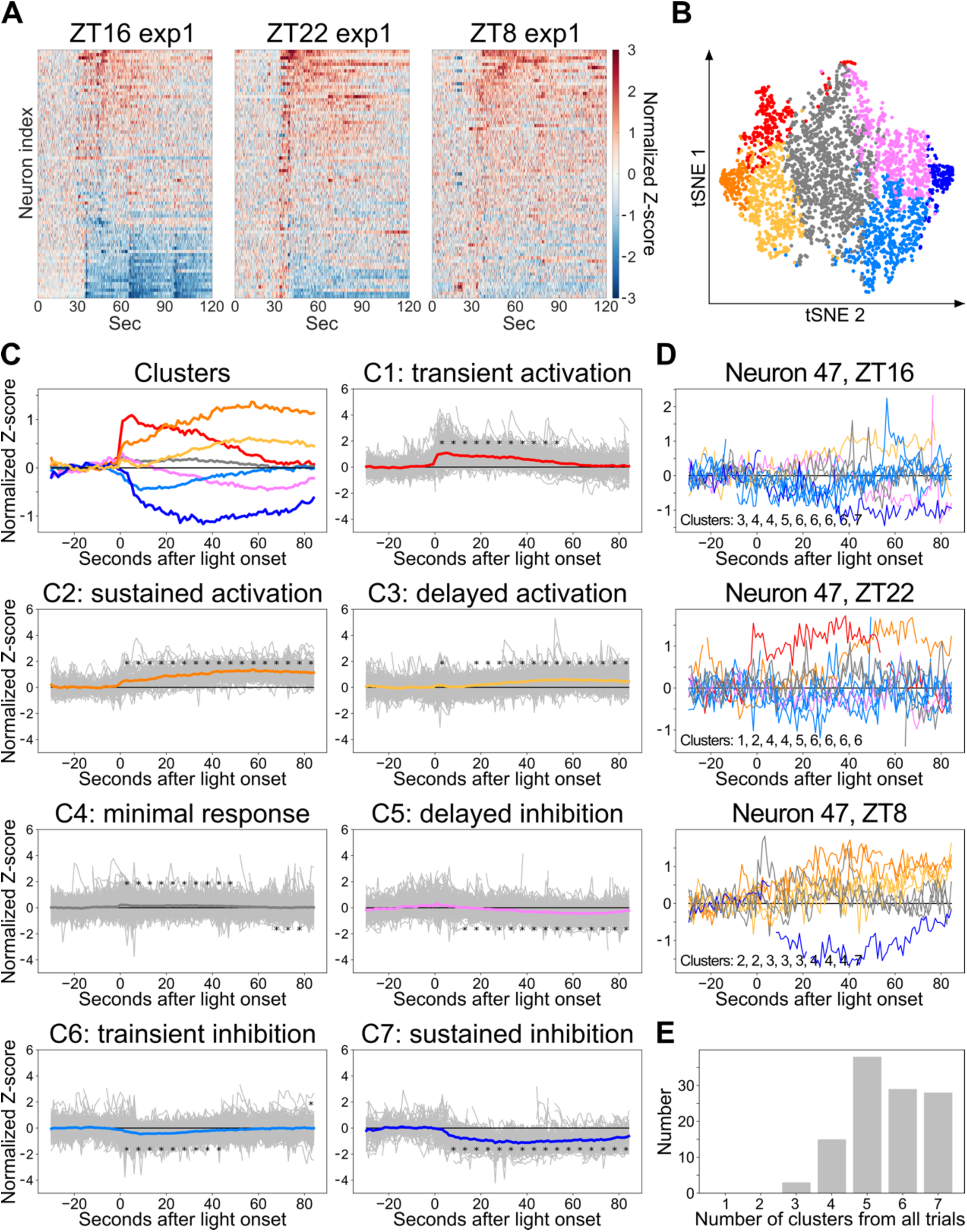
Diverse neuronal light responses in the SCN. **A.** Representative heat map with GCaMP traces (normalized Z-score) from 113 identified neurons at ZT 16 (left), ZT 22 (middle), and ZT 8 (right), respectively. **B-C.** 2D t-SNE map (B) shows the diversity of neuronal light-responses. 2875 recording traces from 113 neurons are plotted, excluding those with excessive motion artifacts. The 2875 recordings are classified into 7 clusters, indicated by different colors, with k-means. The 7 light responses are plotted in (C), including averaged traces of all clusters (upper-left panel) and raw traces (gray) with average from each cluster (the other 7 panels). Stars above or below the average traces indicate significantly higher or lower z-score compared to the -5∼0 second baseline respectively. One-way ANOVA and Tukey post hoc test, * indicates p < 0.05. **D.** Raw traces from the representative neuron that show high diversity of light-evoked responses through 9 repeated trials from 3 time points. **E.** Histogram of neurons shows multiple types of light responses. n = 3 animals.

### Functional Homogeneity in Light Response Patterns of VIP and Non-VIP SCN Neurons

The suprachiasmatic nucleus (SCN) is known to contain a diversity of neurons, including those expressing vasoactive intestinal peptide (VIP) and arginine vasopressin (AVP). Within this traditional molecular classification, VIP^+^ and gastrin-releasing peptide (GRP^+^) neurons have been viewed as the primary light reception unit in the SCN. To investigate the specificity of VIP neurons in light response relative to non-VIP neurons, we next analyzed the difference between tdTomato-positive (VIP^+^) and tdTomato-negative (VIP^-^) neurons. Contrary to expectations, our analysis revealed no significant differences in the average light responses between VIP^+^ and VIP^-^ neurons (Figure 3A), nor in the composition of their light responses (Figure 3B). We further quantified the similarity of light response by computing Pearson correlation coefficients within and between VIP^+^ and VIP^-^ neurons in the same trial (Figure 3C). If VIP neurons constituted a discrete functional cluster within the SCN’s photoentrainment circuitry, one would anticipate a higher in-group correlation coefficient for VIP^+^ neurons. Yet, our results demonstrated that in-group correlations for VIP^+^ neurons did not significantly deviate from correlation coefficients between VIP^+^ and VIP^-^ neurons across all examined time points (Figure 3D). Intriguingly, only during the photoentrainment dead zone at ZT 8 did VIP^+^ neurons exhibit a notably higher correlation than VIP^-^ neurons, suggesting a potential distinct role during this specific phase (Figure 3D, right panel). Further exploration into the contributions of VIP neurons within the dead zone of photoentrainment is required to decipher this phenomenon. Collectively, these findings challenge the canonical view of a distinct VIP neuron-led functional hierarchy for photoentrainment, highlighting a surprising complexity among neuronal light responses within the SCN.

**Figure 3.**
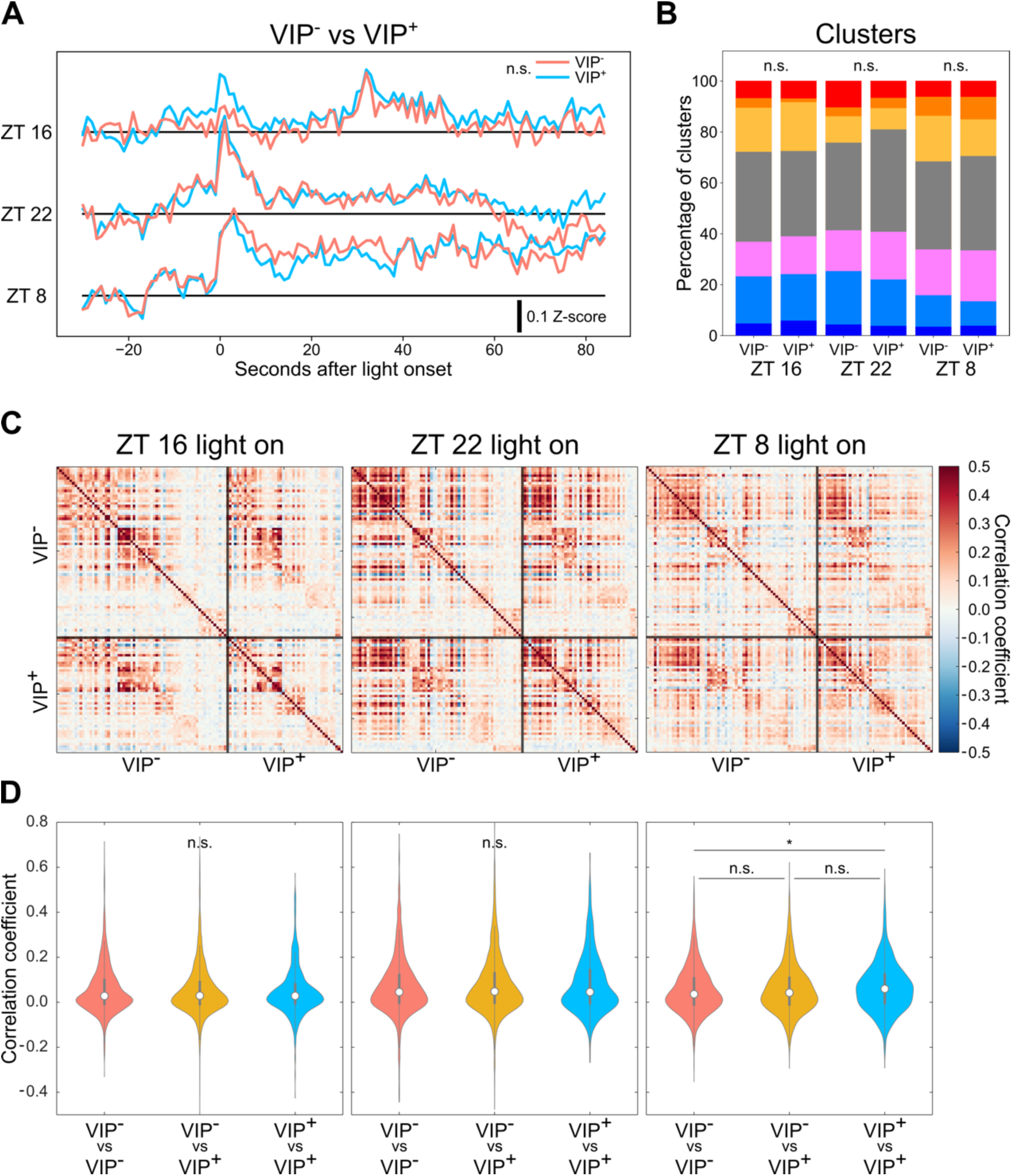
VIPergic neurons show similar light responses to non-VIP neurons. **A.** Averaged light response traces from VIP^+^ neurons (blue) and VIP^-^ neurons (pink). n.s. indicates p > 0.05, two-way ANOVA, Tukey post hoc tests. **B.** Composition of 7 clusters of light response at different time points among VIP^+^ and VIP^-^ neurons. No significantly difference is observed between VIP^+^ and VIP^-^ neurons (two-way ANOVA). **C.** Correlation maps comparing VIP^+^ and VIP^-^ neurons. Each pixel on the map represents the Pearson correlation coefficient between two neurons. **D.** Violin plots summarize the Pearson correlation coefficient of each comparison pair among cell types, including VIP^-^ to VIP^-^ (pink), VIP^+^ to VIP^-^ (yellow), and VIP^+^ to VIP^+^ (blue). The star indicates p < 0.05 from one-way ANOVA, Tukey post hoc tests. n = 3 animals. For violin plot, circle indicates median, think vertical line indicates interquartile range and thin vertical line indicates 1.5X interquartile range.

### Time-Dependent Variation in SCN Neuronal Responses to Light

Circadian photoentrainment is characterized by three discrete light responses. Here we analyzed the recording of 113 neurons to dissect the light response dynamics of SCN neurons at a single-cell level across these temporal zones. Our comparative analysis of seven response clusters at different ZTs revealed no significant differences in most clusters at the population level (Extended Figure 3A). Notably, the proportion of neurons exhibiting an inhibitory response (clusters 6 or combined cluster 6 with 7) was significantly reduced at ZT 8 (Figure 4A), reflecting the subtle differences in population-wide light response patterns (Figure 2A and Extended Figure 2). When the heatmap was organized in the same order as ZT 16, stark contrasts in individual neuronal responses emerged across the three time points (Figure 4B). To investigate the potential differences in SCN’s time-gated network properties, we calculated the Pearson correlation coefficient of fluorescence intensities between neurons across all recorded datasets. If SCN is comprised with a stable, time-independent network, one would be expected to observe consistent light responses across different ZTs for individual neurons, which will generate a high correlation coefficient. Surprisingly, our cross-time correlation analysis during the 90-second light exposure revealed correlation coefficients close to zero (R-cross, ZT 8 – ZT 16, ZT 16 – ZT 22, and ZT8 – ZT 22, etc.) (Figure 4C blue outline, Figure 4D). Additionally, the correlation coefficients for different neurons at the same ZT time within a single trial (R-same, ZT 8, ZT 16, ZT 22) (Figure 4C yellow outline) were significantly higher than the R-Cross values (Figure 4D). These findings indicate that SCN neurons show some degree of synchronization within each trial. However, light responses for individual neuronal vary with different trials and ZT, suggesting a dynamic functional network governing circadian photoentrainment.

**Figure 4.**
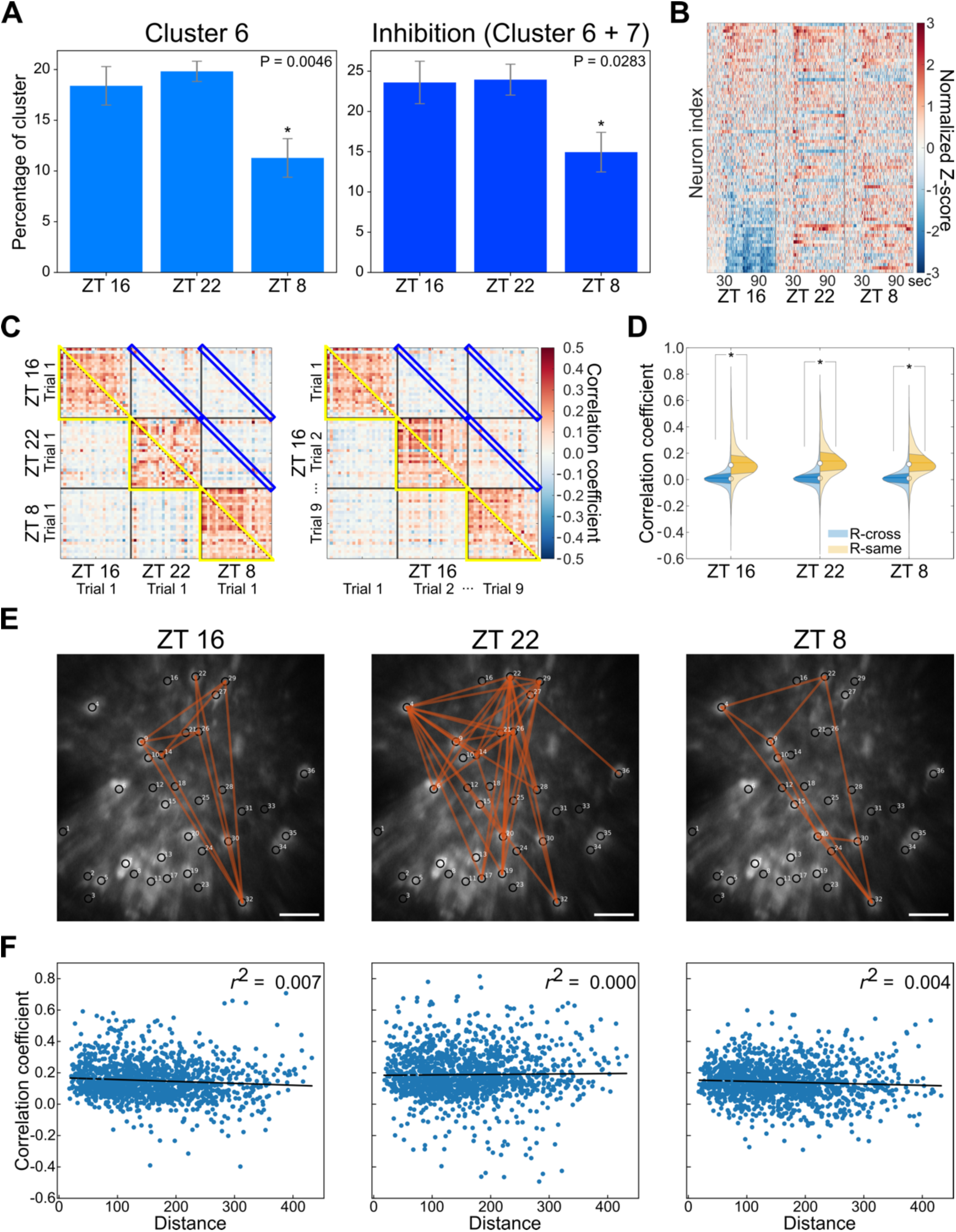
Dynamic light responses from individual SCN neurons that form distinct temporal circuit. **A.** Percentage of inhibitory clusters is significantly decreased at ZT 8. **B.** Heat map of combined normalized traces from three time points for 113 identified neurons ranked by mean Z-score at ZT16. **C.** Representative heat maps of Pearson correlation coefficients comparing light responses between neurons. Left panel shows comparison between different ZTs, right panel shows comparison between different trials within the same ZT. **D.** Cross-time correlation analysis from each neuron between different trials (R-cross, blue outline in C and blue plots in D) is significantly lower than the correlation coefficients for different neurons at the same ZT time within a single trial (R-same, yellow outline in C and orange plots in D). **E.** Representative diagram showing highly correlated neuron pairs (average r from 9 repeats > 0.5) from each time points. **F.** Scatter plots of distance between pairs of neurons and their Pearson correlation coefficient in respective time points. The r^2^ value of linear regression lines is 0.007, 0, and 0.004 for ZT 16, ZT 22, and ZT 8 respectively. For (A) and (D), * indicates p < 0.05 with one-way ANOVA and Tukey post hoc tests. Scale bars are 100 μm, according to the imaging side of the GRIN endoscope. n = 3 animals. For violin plot, circle indicates median, think vertical line indicates interquartile range and thin vertical line indicates 1.5X interquartile range.

To determine whether this network is driven by local connections, we identified neurons with high connectivity based on their correlation coefficients (r > 0.5). We found three distinct functional connectivity maps corresponding to ZT 8, ZT 16, and ZT 22, with no clear spatial clustering among the highly connected neurons (Figure 4E and Extended Figure 4). We then mapped the correlation coefficients against the inter-neuronal distances during the light exposure period across all three time points. A locally driven network would typically exhibit a negative correlation between neuronal distance and correlation coefficient. Contrary to this expectation, the relationship between the neurons’ connectivity and their physical proximity was negligible, with an average r-squared value of only 0.04 (Figure 4F). Collectively, these observations suggest that SCN neuronal light responses are orchestrated by a complex network architecture rather than simple local circuits.

### Identification of Three Distinct SCN Neuronal Groups and Their Roles in Phase Shifting and Circadian Computation

To determine if the SCN consists of neuron populations with time-dependent light responses, we conducted additional analyses combining light responses from three distinct time points of each neuron. First, we calculated the average fluorescence z-score for individual neurons in five-second bins to reduce the effect of transient calcium waves. Subsequent ANOVA tests compared 18 light exposure bins to baseline across nine trials for each neuron. Bins showing significant differences in post-hoc tests (Figure 5A) were marked in orange to indicate significantly higher fluorescence intensity, and in blue for significantly lower intensity compared to their respective baselines (Figure 5B). While most neurons exhibited varying light responses in each trial, only a small population showed consistently significant positive or negative responses. Through k-means clustering of data from all 54 bins across the three time points, we identified three distinct groups of neurons (Figure 5B). Specifically, one group displayed a persistent positive light response at ZT16 (group 1, 11.5%), another showed consistent inhibition only at ZT22 (group 2, 4.4%), and the majority did not exhibit a consistent light response (group 3, 84.1%), resulting in an overall flat response curve (Figure 5C). When we compare the light response composition, group 1 neurons showed higher percentage of activation clusters (cluster 1-4) at ZT 16, group 2 neurons showed higher percentage of inhibition clusters (cluster 5+7) at ZT 22 (Figure 5D). Next, we generated time-correlated Haar features for each neuron by analyzing the polarity of 18 bins from each time point, revealing distinct patterns for each cluster in the Haar-ZT space. Interestingly, group 1 was differentiated from groups 2 and 3 at ZT16, whereas group 2 was distinct from groups 1 and 3 at ZT22. No clear segregation was observed among the three groups at ZT8 (Figure 5E and 5F). Interestingly, both VIP^+^ and VIP^-^ neurons were present in groups 1 and 2, with neurons in group 1 exclusively observed in the ventral focus planes of our recordings (Figure 5G). It is noteworthy that most group 3 neurons (72.6%) still displayed a range of light responses throughout the nine trials at specific time point. The diversity of their light response types was significantly higher than that observed in baseline (Extended Figure 5). Collectively, these results demonstrate that the SCN’s functional network operates on a time-gated mechanism. Moreover, we have identified two subpopulations of SCN neurons with consistent light responses at ZT 16 or ZT 22 respectively, while the majority show dynamic light responses between trials.

**Figure 5.**
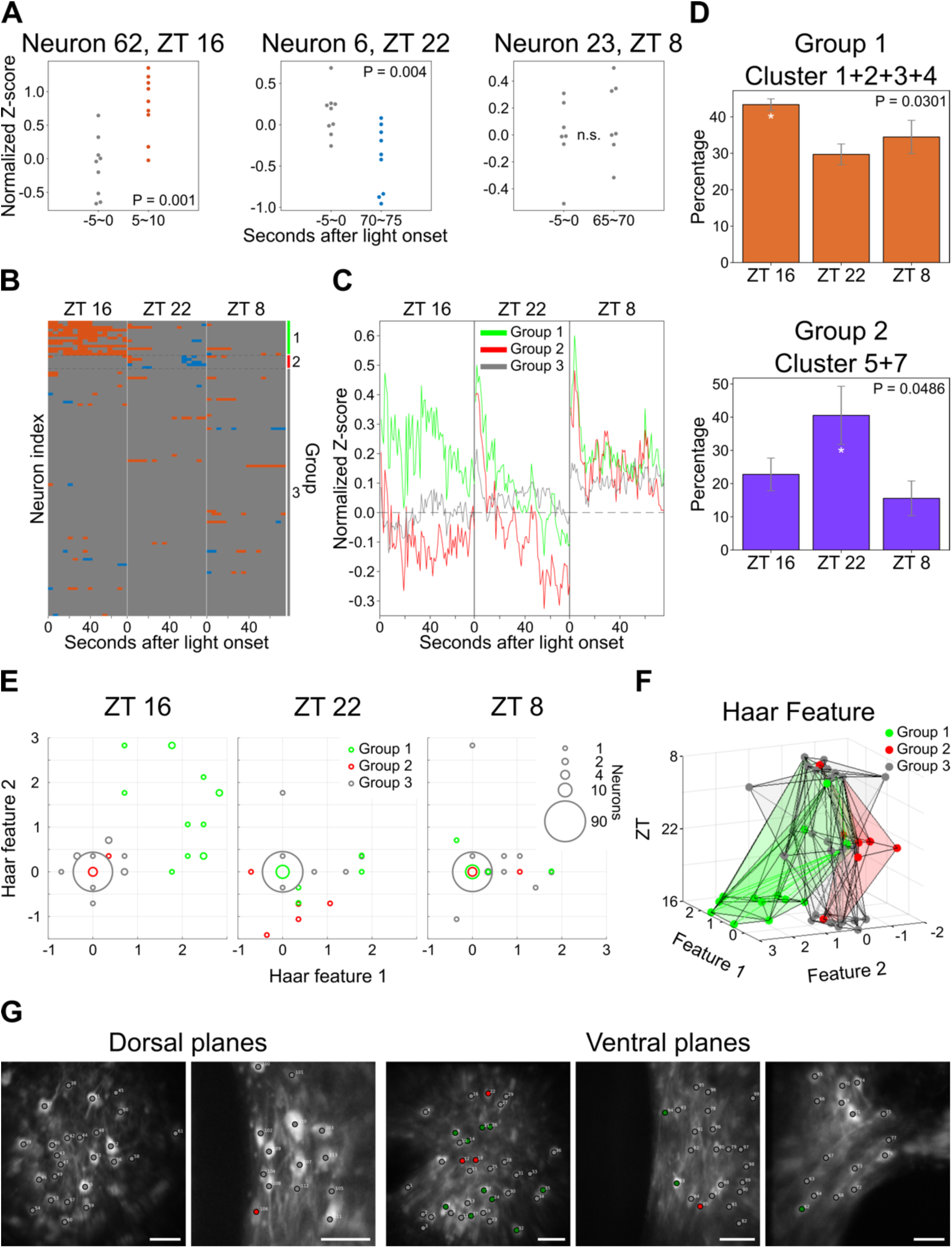
Three distinct groups of SCN neurons with different time dependent light responses. **A.** Representative light response analysis from 3 neurons. Three possible results are demonstrated here, including that the Z-score of the specific time point is significantly higher (orange), lower (blue), or not significantly changed (grey) compared to the baseline (one-way ANOVA and Tukey test, p < 0.05). **B.** The results in A are plotted together. Neurons are classified into three groups according to their significance pattern across all time points, separated by dotted lines. Group 1 neurons (green) are specific activated at ZT 16. Group 2 neurons (red) are specifically inhibited at ZT 22. While group 3 neurons (grey) do not show consistent positive or negative light response. **C.** Averaged light-response traces from three groups of neurons at different time points. **D.** Group 1 neurons show significantly higher activation clusters (cluster 1-4) at ZT 16 and group 2 neurons show significantly higher inhibition clusters (5 and 7) at ZT 22. * indicates p < 0.05 with one-way ANOVA and Tukey post hoc tests. **E.** Haar feature analysis of all neurons for each ZT using 18 bins of 5 sec average fluorescence intensity. **F.** Neuron trajectories in the Haar-ZT space show distinct characters from three groups. **G.** Neuron groups are labeled on the raw images. Green dots indicate the cell body of group 1 neurons, red dots indicate group 2 neurons, while grey dots indicate group 3 neurons. Scale bars are 100 μm, according to the imaging side of the GRIN endoscope. n = 3 animals.

### Activation of CT 16-trapped light-responsive neurons produces phase delay and breaks circadian photoentrainment dead zone

Since a particular group of neurons consistently exhibits a positive light response at ZT 16, we asked whether activation of these neurons could induce phase delays at any time of the circadian cycle. To test whether SCN neurons utilize specific neuronal circuitry for discrete circadian photoentrainment, we used TRAP2 mice with Cre-dependent AAV containing excitatory Gs-coupled DREADDs (rM3Ds) to target neurons within the SCN that are responsive to light at CT 16 or CT 22 (Figure 6A). Four weeks post-injection with either AAV9-hSyn-DIO-rM3D(Gs)-mCherry or AAV9-hSyn-DIO-EGFP into the SCN, we randomly divided the mice into two groups. Each group was subjected to a 900 lux, 10-minute light pulse at CT 16 or CT 22, followed by an injection of 4-OHT to label light-responsive SCN neurons as TRAP-CT 16 or TRAP-CT 22 (Figure 6B and Extended Figure 6). We then monitored wheel-running activity in constant darkness and administered CNO or PBS as a control at various times to assess whether the activation of TRAP-CT 16 or TRAP-CT 22 neurons could mimic the phase shifts typically induced by light (Figure 6C-E and Extended Figure 7). Given that CNO is known to affect DREADD neurons for 4-6 hours post-injection, we strategically administered it at CT 2 to target the dead zone. Remarkably, the activation of TRAP-CT 16 neurons led to a significant phase delay across all three time points, including the photoentrainment dead zone and advance zone, in contrast to control mice injected with CNO (Figure 6F). Notably, activation of TRAP-CT 22 neurons resulted in a significant phase delay at CT 16, a significant phase advance at CT 22, and no phase shift at CT 2, similar to the usual light-induced phase response curve (Figure 6G). This result suggested a dynamic functional circuit that activation of CT 22-trapped neurons could produce multiple behavior responses at specific circadian time. Together, our findings indicate the presence of a bi-stable circuit that can lead to at least two distinct time-gated functional networks within the SCN for circadian photoentrainment. Furthermore, the output sub-population responsible for phase delays in the SCN can be isolated.

**Figure 6.**
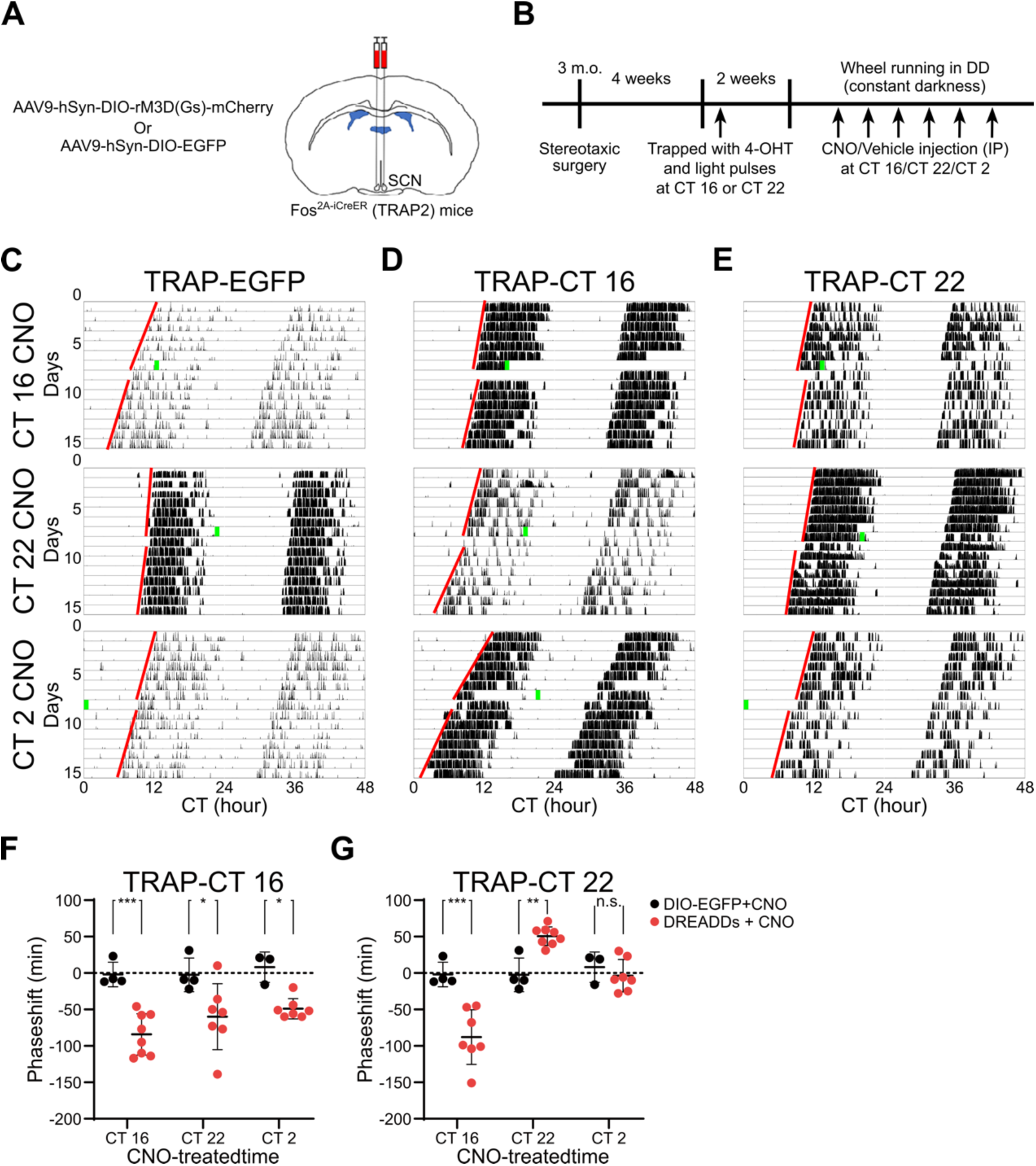
Activation of CT 16-trapped light-responsive neurons produces phase delay and breaks the circadian photoentrainment dead zone. **A.** Experimental scheme for DREADDs bilateral SCN injection in the TRAP2 (Fos-iCreER) mice. **B.** Time table for experimental procedure. **C.** Representative actogram for GFP-expressing CT 16 or CT 22-trapped mice before and after CNO injection. **D.** Representative actogram for DREADDs (rM3Ds)-expressing CT 16-trapped mice before and after CNO injection. **E.** Representative behavioral data for DREADDs (rM3Ds)-expressing CT 22-trapped mice before and after CNO injection. Green bars indicate the time points of CNO injection, while red lines indicate linear regression line of activity onsets. **F.** Statistics of phase shift analysis for GFP-expressing, DREADDs (rM3Ds)-expressing CT 16-trapped, and CT 22-trapped mice. Two-way ANOVA: * indicates p < 0.05, ** indicates p < 0.01, *** indicates p < 0.001, n = 3-6 for control group, n = 7-8 for CT 16-trapped group, n = 6-7 for CT 22-trapped group.

**Figure 7.**
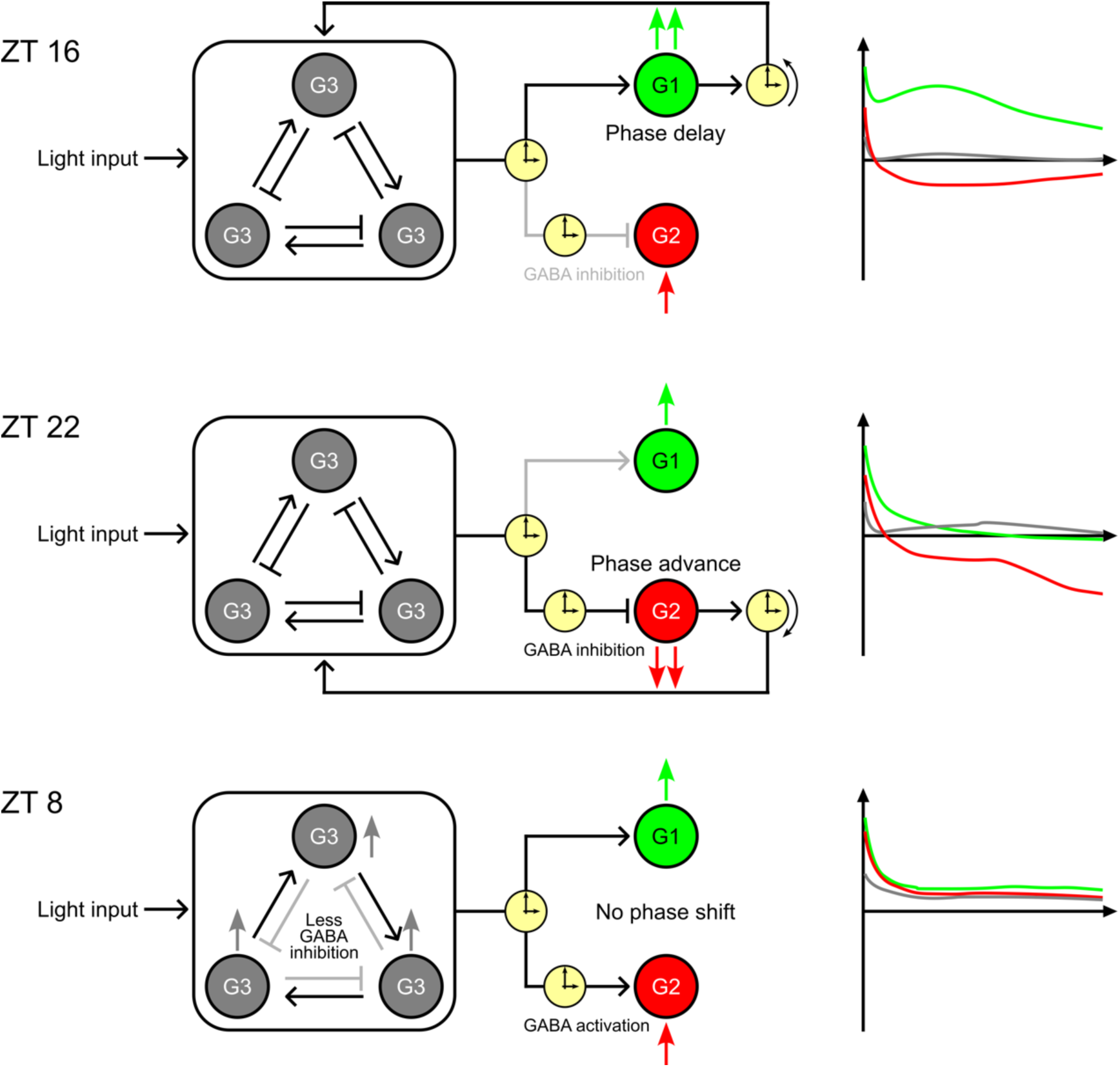
Hypothetic model of SCN circuit for circadian photoentrainment. Proposed bi-stable functional circuit within the SCN comprised with 3 distinct groups of neurons. Group 1 and 2 neurons are output units to drive phase shift at delay or advance time respectively. Group 3 neurons are the computational unit using populational dynamic coding.

## Discussion

In this paper, we demonstrate the circadian time-dependent heterogeneity of SCN neurons in response to acute light, indicating a dynamic network within the SCN for circadian phase responses. Despite decades of study, several physical barriers have hindered the direct observation of the SCN in living animals. Located deep within the brain and hypothalamus, nearly 6 mm from the skull surface in mice, the SCN’s accessibility to most optical research approaches is obstructed. Additionally, the SCN, being a small nucleus of 400 micrometers or less in every dimension and consisting of approximately 20,000 neurons, limits the feasibility of electrical recording.

Consequently, *ex vivo* SCN preparations on brain slides have predominated as the methodology for studying mammals’ central clock. However, this approach severs the SCN’s endogenous network from RHT input and other brain regions, potentially altering the master clock’s innate properties, including photoentrainment functions. Through the use of *in vivo* calcium imaging combined with a two-photon microscope and GRIN endoscopes, we have identified several emergent properties of the SCN. These include immediate responses to acute light stimulation, circadian time-dependent variable light response patterns, and neurons that are selectively responsive to light at specific circadian times. These emergent properties highlight the necessity of an *in vivo* setup with minimal network interruption to understand the central clock’s computational network, which shall also be beneficial for studying other brain nuclei.

Our findings suggest an unconventional information flow in the SCN to process ambient light information, which was described as a simple linear circuit from the RHT to the VIP and GRP neurons-enriched ventrolateral “core”, and then to the AVP neurons enriched dorsomedial “shell” region. In this prevalent structure, the PRC has to be integrated by a specific subset of neurons in SCN that are constantly responsive to light input. In other words, a certain subset of neurons in the SCN would show consistent light responses in all three circadian time zones. However, our findings show that distinct neuron groups are respectively responsive to light at different circadian time zones. Combining with recent anatomical studies showing dense ipRGC axonal fibers in the SCN ^17–19,72^, our results indicating that a complex network structure including many feedback circuits in SCN may better describe circadian photoentrainment than the conventional simple hierarchical model.

In our study, we have shown that neurons responsive to acute light in the SCN can be categorized by their timing of light response. We identified distinct neuronal groups that respond specifically to light during the time zones associated with phase delays or advances. This suggests that the bidirectionality of the PRC consists of at least two distinct functional circuits within the SCN: neurons activated during early subjective night that contribute to phase delays, and neurons inhibited during late subjective night that contribute to phase advances. Our findings are in line with previous research identifying a spatiotemporal light responsive pattern in the SCN ^30,40^. Behaviorally, we demonstrated that activation of CT 16-trapped neurons leads to phase delays across all time zones, supporting the idea of separable delay and advance circuitries within the SCN. Prior research indicated a partial PRC by activating VIP or CCK neurons ^36,56^, hinting that a subset of these may correspond to our identified delay or advance cells. Therefore, these neurons may be output neurons which contribute higher weight to synchronize the whole SCN population for phase shift. However, due to the absence of specific genetic tools for trapping inhibited neurons, we could not directly confirm the causal relationship between ZT 22 inhibition and the behavioral phase advances. We hypothesize that a portion of what may be group 3 neurons, involved in the population coding for light input, were captured in TRAP-CT 22 mice. Consequently, the activation of these CT 22-trapped neurons generates similar effect to light exposure. This explains why their activation does not solely induce phase advances but can also cause typical light-induced phase delays or no phase shift, depending on the time of activation, suggesting a bi-stable outcome for phase responses. Moreover, the independent modulation of circadian phase shifts by two groups of neurons suggests a potential role in encoding daylength and seasonal rhythms, which are not yet well understood in mammals.

Combining our results, we propose the existence of two ’time-gates’ within the SCN’s dynamic network. The first ’time-gate’ operates similarly to a bi-stable transcriptional network model seen in cell fate differentiation. This gate could create two distinct low-entropy states within the SCN: one that leads to the activation of group 1 neurons, facilitating phase delay, and another that results in the inhibition of group 2 neurons, leading to phase advance. These low-entropy states reflect the dichotomous nature of the network’s response to light, reminiscent of the robust and divergent outcomes observed in cellular differentiation pathways. The second proposed ’time-gate’ in the SCN’s dynamic network is characterized by a reduction in inhibition at ZT 8, potentially disrupting the feedback and feedforward interactions that are critical for the bi-stable model’s function. This disturbance could prevent the network from settling into either of the two low-entropy states associated with phase shifts, offering an explanation for the absence of behavioral phase shifts after light exposure during the dead zone. An interesting analogy is the bi-stability in electronics. In a typical electronic bistable circuit, the basic components are feedback and feedforward between two inverters, which form a logic unit. Our findings of reduced inhibition during ZT 8 also in line with previous research showing a diurnal switch in GABAergic signaling within the SCN, where it transitions from inhibitory to excitatory during the mid-day ^73,74^. These results substantiate our proposed bi-stable network model and supports the validity of our observations and theoretical framework for circadian photoentrainment in the SCN (Figure 7).

Surprisingly, we also found that most neurons exhibited dynamic light response types across trials. The observed stochastic response of neurons to light within the SCN population, despite the seemingly straightforward process of phase shifting, suggests a highly flexible network. Recent studies have proposed a dynamic population coding in the cortex ^75^. Here we observed similar feature in the SCN circuit. Therefore, this dynamics network property may be a universal feature of the mammalian central nervous system that is integrated to the SCN for photoentrainment computation. This flexibility allows for a large pool of neurons to be available for input and computation, with subsets being utilized in a seemingly random fashion for each trial. This strategy preserves the system’s plasticity for learning process, and offers redundancy to safeguard against damage. Such a mechanism ensures that even if individual neurons are compromised, the network retains its computational capabilities, channeling the processing through a select group of outcome-determining neurons.

The methodology and analysis carried out from current study also has their limitations. One limitation of our study is the slower temporal resolution of calcium imaging compared to electrophysiological recording techniques. While calcium imaging is effective for observing global light responses over relatively long timescales, it does not have the temporal resolution to resolve the difference between monosynaptic or polysynaptic responses, or detecting spike in SCN neurons. Our technique primarily captures the overall light response of the SCN network, potentially obscuring more subtle, rapid synaptic events. Consequently, the similar responses we observed from VIP^+^ and VIP^-^ neurons cannot be conclusively interpreted as evidence of similar synaptic connections from the ipRGCs. Furthermore, we could not differentiate whether the inhibition light response we observed is the result of di-synaptic input through SCN GABAergic signal or direct GABA signal from ipRGCs ^76^. Another limitation arises from our focus on analyzing activity trends during 90 seconds of light exposure, rather than the oscillation of calcium transients observed in many SCN neurons. Previous research indicates that the frequency of spontaneous calcium transient events varies with the time of day *in vivo* ^35^. By concentrating on the general light response across 27 trials, we have not characterized these oscillations due to their phase could not be aligned perfectly to our recording paradigm. Since the physiological function and origin of spontaneous calcium transient in the SCN neurons is still unclear. Our approach may overlook the implications of calcium transient for understanding the complete picture of circadian photoentrainment and SCN functionality. Finally, we recorded the calcium-mediated light responses in the SCN at two different Z positions, approximately 105 µm apart. Although the dorsal section, on average, contains fewer VIP^+^ neurons, we cannot conclusively determine whether the ventral or dorsal Z sections correspond to the core or shell regions of the SCN, respectively. It remains possible that both Z sections are situated within either the core or the shell region. The long GRIN endoscopes required for accessing and recording from SCN neurons possess high pitch and low numerical aperture (NA). This limitation restricts the excitation efficiency of GCaMP within the two-photon system, thereby constraining our ability to achieve clear recordings when attempting to span larger Z-axis distances between two focal planes. In the future, a new, higher NA rod-shaped thin endoscopes with a diameter of 600 µm or less, which can be consistently inserted above the SCN, will be required to overcome this challenge.

In summary, our methodological approach, employing two-photon excitation for spatial analysis, offers a minimally perturbative tool for studying SCN networks longitudinally. This advancement opens doors to exploring the fine-grained spatial organization of the SCN at single cell level *in vivo* and its implications for circadian photoentrainment. Our study unveils the intricate network nature of the mammalian circadian clock within the SCN. The identification of distinct neuronal groups that are responsive to light at different circadian times provides new insights beyond the traditional hierarchical model of circadian photoentrainment. This discovery lays the groundwork for future research into the identification of molecular markers and functional roles of the SCN neuronal subtypes, illuminating the underlying mechanisms for matching the endogenous clock with external light-dark cycle in mammals.

## Acknowledgements

This work was supported by the Taiwan National Science and Technology Council grant NTSC 112-2636-B-002-012, 112-2628-B-002-028, and 112-2321-B-002 -019 (to S.-K.C.) and National Taiwan University. We thank the Technology Commons, College of Life Science at National Taiwan University for technical assistance with confocal imaging and virus core for virus packing. We also thank Yin-Tzu Hsieh for generating the illustration in figures.

## Data availability

This study includes no data deposited in external repositories.

## Reference

1 Ono, D. et al. The Suprachiasmatic Nucleus at 50: Looking Back, Then Looking Forward. Journal of biological rhythms 39, 135–165 (2024). 10.1177/07487304231225706

2 Hirota, T. & Fukada, Y. Resetting mechanism of central and peripheral circadian clocks in mammals. Zoolog Sci 21, 359–368 (2004). 10.2108/zsj.21.359

3 Yamazaki, S. et al. Resetting central and peripheral circadian oscillators in transgenic rats. Science 288, 682–685 (2000).

4 Ralph, M. R., Foster, R. G., Davis, F. C. & Menaker, M. Transplanted suprachiasmatic nucleus determines circadian period. Science 247, 975–978 (1990). 10.1126/science.2305266

5 Hastings, M. H., Maywood, E. S. & Brancaccio, M. Generation of circadian rhythms in the suprachiasmatic nucleus. Nat Rev Neurosci 19, 453–469 (2018). 10.1038/s41583-018-0026-z

6 Hattar, S. et al. Melanopsin and rod-cone photoreceptive systems account for all major accessory visual functions in mice. Nature 424, 76–81 (2003). 10.1038/nature01761 nature01761 [pii]

7 Provencio, I., Rollag, M. D. & Castrucci, A. M. Photoreceptive net in the mammalian retina. This mesh of cells may explain how some blind mice can still tell day from night. Nature 415, 493 (2002).

8 Panda, S. et al. Melanopsin (Opn4) requirement for normal light-induced circadian phase shifting. Science 298, 2213–2216 (2002). 10.1126/science.1076848 298/5601/2213 [pii]

9 Hattar, S., Liao, H. W., Takao, M., Berson, D. M. & Yau, K. W. Melanopsin-containing retinal ganglion cells: architecture, projections, and intrinsic photosensitivity. Science 295, 1065–1070 (2002). 10.1126/science.1069609 295/5557/1065 [pii]

10 Berson, D. M., Dunn, F. A. & Takao, M. Phototransduction by retinal ganglion cells that set the circadian clock. Science 295, 1070–1073 (2002).

11 Gooley, J. J., Lu, J., Chou, T. C., Scammell, T. E. & Saper, C. B. Melanopsin in cells of origin of the retinohypothalamic tract. Nat Neurosci 4, 1165 (2001).

12 Provencio, I. et al. A novel human opsin in the inner retina. J Neurosci 20, 600–605 (2000).

13 Panda, S. et al. Melanopsin is required for non-image-forming photic responses in blind mice. Science 301, 525–527 (2003). 10.1126/science.1086179

14 Johnson, R. F., Moore, R. Y. & Morin, L. P. Loss of entrainment and anatomical plasticity after lesions of the hamster retinohypothalamic tract. Brain Res 460, 297–313 (1988). https://doi.org/0006-8993(88)90374-5 [pii]

15 Chew, K. S. et al. A subset of ipRGCs regulates both maturation of the circadian clock and segregation of retinogeniculate projections in mice. Elife 6 (2017). 10.7554/eLife.22861

16 Fernandez, D. C., Chang, Y. T., Hattar, S. & Chen, S. K. Architecture of retinal projections to the central circadian pacemaker. Proc Natl Acad Sci U S A 113, 6047–6052 (2016). 10.1073/pnas.1523629113

17 Hattar, S. et al. Central projections of melanopsin-expressing retinal ganglion cells in the mouse. J Comp Neurol 497, 326–349 (2006). 10.1002/cne.20970

18 Chen, S. K., Badea, T. C. & Hattar, S. Photoentrainment and pupillary light reflex are mediated by distinct populations of ipRGCs. Nature 476, 92–95 (2011). 10.1038/nature10206

19 Ecker, J. L. et al. Melanopsin-Expressing Retinal Ganglion-Cell Photoreceptors: Cellular Diversity and Role in Pattern Vision. Neuron 67, 49–60 (2010). https://doi.org/S0896-6273(10)00419-8 [pii] 10.1016/j.neuron.2010.05.023

20 Hatori, M. et al. Inducible ablation of melanopsin-expressing retinal ganglion cells reveals their central role in non-image forming visual responses. PLoS ONE 3, e2451 (2008).

21 Guler, A. D. et al. Melanopsin cells are the principal conduits for rod-cone input to non-image-forming vision. Nature 453, 102–105 (2008). https://doi.org/nature06829 [pii] 10.1038/nature06829

22 Khalsa, S. B., Jewett, M. E., Cajochen, C. & Czeisler, C. A. A phase response curve to single bright light pulses in human subjects. J Physiol 549, 945–952 (2003). 10.1113/jphysiol.2003.040477

23 Pittendrigh, C. S. Circadian rhythms and the circadian organization of living systems. Cold Spring Harb Symp Quant Biol 25, 159–184 (1960). 10.1101/sqb.1960.025.01.015

24 Daan, S. & Pittendrigh, C. S. A functional analysis of circadian pacemakers in nocturnal rodents. II. The variability of phase response curves. Journal of Comparative Physiology A 106, 253–266 (1976).

25 Foster, R. G., Hughes, S. & Peirson, S. N. Circadian Photoentrainment in Mice and Humans. Biology (Basel) 9 (2020). 10.3390/biology9070180

26 Ni, J. D., Baik, L. S., Holmes, T. C. & Montell, C. A rhodopsin in the brain functions in circadian photoentrainment in Drosophila. Nature 545, 340–344 (2017). 10.1038/nature22325

27 Emery, P. et al. Drosophila CRY is a deep brain circadian photoreceptor. Neuron 26, 493–504 (2000). 10.1016/s0896-6273(00)81181-2

28 Stanewsky, R. et al. The cryb mutation identifies cryptochrome as a circadian photoreceptor in Drosophila. Cell 95, 681–692 (1998). 10.1016/s0092-8674(00)81638-4

29 Shigeyoshi, Y. et al. Light-induced resetting of a mammalian circadian clock is associated with rapid induction of the mPer1 transcript. Cell 91, 1043–1053 (1997). 10.1016/s0092-8674(00)80494-8

30 Yan, L. & Silver, R. Differential induction and localization of mPer1 and mPer2 during advancing and delaying phase shifts. Eur J Neurosci 16, 1531–1540 (2002). 10.1046/j.1460-9568.2002.02224.x

31 Albrecht, U., Sun, Z. S., Eichele, G. & Lee, C. C. A differential response of two putative mammalian circadian regulators, mper1 and mper2, to light. Cell 91, 1055–1064 (1997). 10.1016/s0092-8674(00)80495-x

32 Xu, P. et al. NPAS4 regulates the transcriptional response of the suprachiasmatic nucleus to light and circadian behavior. Neuron 109, 3268–3282 e3266 (2021). 10.1016/j.neuron.2021.07.026

33 Nielsen, H. S., Hannibal, J., Knudsen, S. M. & Fahrenkrug, J. Pituitary adenylate cyclase-activating polypeptide induces period1 and period2 gene expression in the rat suprachiasmatic nucleus during late night. Neuroscience 103, 433–441 (2001). 10.1016/s0306-4522(00)00563-7

34 Ding, J. M. et al. Resetting the biological clock: mediation of nocturnal circadian shifts by glutamate and NO. Science 266, 1713–1717 (1994). 10.1126/science.7527589

35 Irwin, R. P. & Allen, C. N. Calcium response to retinohypothalamic tract synaptic transmission in suprachiasmatic nucleus neurons. J Neurosci 27, 11748–11757 (2007). 10.1523/JNEUROSCI.1840-07.2007

36 Jones, J. R., Simon, T., Lones, L. & Herzog, E. D. SCN VIP Neurons Are Essential for Normal Light-Mediated Resetting of the Circadian System. J Neurosci 38, 7986–7995 (2018). 10.1523/JNEUROSCI.1322-18.2018

37 Mazuski, C. et al. Entrainment of Circadian Rhythms Depends on Firing Rates and Neuropeptide Release of VIP SCN Neurons. Neuron 99, 555–563 e555 (2018). 10.1016/j.neuron.2018.06.029

38 Azzi, A. et al. Network Dynamics Mediate Circadian Clock Plasticity. Neuron 93, 441–450 (2017). 10.1016/j.neuron.2016.12.022

39 Azzi, A. et al. Circadian behavior is light-reprogrammed by plastic DNA methylation. Nat Neurosci 17, 377–382 (2014). 10.1038/nn.3651

40 Duy, P. Q. et al. Light Has Diverse Spatiotemporal Molecular Changes in the Mouse Suprachiasmatic Nucleus. Journal of biological rhythms 35, 576–587 (2020). 10.1177/0748730420961214

41 Evans, J. A., Leise, T. L., Castanon-Cervantes, O. & Davidson, A. J. Dynamic interactions mediated by nonredundant signaling mechanisms couple circadian clock neurons. Neuron 80, 973–983 (2013). 10.1016/j.neuron.2013.08.022

42 Myung, J. & Pauls, S. D. Encoding seasonal information in a two-oscillator model of the multi-oscillator circadian clock. Eur J Neurosci 48, 2718–2727 (2018). 10.1111/ejn.13697

43 Myung, J. et al. GABA-mediated repulsive coupling between circadian clock neurons in the SCN encodes seasonal time. Proc Natl Acad Sci U S A 112, E3920–3929 (2015). 10.1073/pnas.1421200112

44 Uriu, K. & Tei, H. Complementary phase responses via functional differentiation of dual negative feedback loops. PLoS Comput Biol 17, e1008774 (2021). 10.1371/journal.pcbi.1008774

45 Varadarajan, S. et al. Connectome of the Suprachiasmatic Nucleus: New Evidence of the Core-Shell Relationship. eNeuro 5 (2018). 10.1523/ENEURO.0205-18.2018

46 Morris, E. L. et al. Single-cell transcriptomics of suprachiasmatic nuclei reveal a Prokineticin-driven circadian network. EMBO J 40, e108614 (2021). 10.15252/embj.2021108614

47 Wen, S. et al. Spatiotemporal single-cell analysis of gene expression in the mouse suprachiasmatic nucleus. Nat Neurosci 23, 456–467 (2020). 10.1038/s41593-020-0586-x

48 Abrahamson, E. E. & Moore, R. Y. Suprachiasmatic nucleus in the mouse: retinal innervation, intrinsic organization and efferent projections. Brain Res 916, 172–191 (2001). 10.1016/s0006-8993(01)02890-6

49 Silver, R. et al. Calbindin-D28K cells in the hamster SCN express light-induced Fos. Neuroreport 7, 1224–1228 (1996). 10.1097/00001756-199604260-00026

50 Romijn, H. J., Sluiter, A. A., Pool, C. W., Wortel, J. & Buijs, R. M. Differences in colocalization between Fos and PHI, GRP, VIP and VP in neurons of the rat suprachiasmatic nucleus after a light stimulus during the phase delay versus the phase advance period of the night. J Comp Neurol 372, 1–8 (1996). 10.1002/(SICI)1096-9861(19960812)372:1<1::AID-CNE1>3.0.CO;2-7

51 Collins, B. et al. Circadian VIPergic Neurons of the Suprachiasmatic Nuclei Sculpt the Sleep-Wake Cycle. Neuron 108, 486–499 e485 (2020). 10.1016/j.neuron.2020.08.001

52 Albers, H. E., Gillespie, C. F., Babagbemi, T. O. & Huhman, K. L. Analysis of the phase shifting effects of gastrin releasing peptide when microinjected into the suprachiasmatic region. Neurosci Lett 191, 63–66 (1995). 10.1016/0304-3940(95)11559-1

53 Reed, H. E., Meyer-Spasche, A., Cutler, D. J., Coen, C. W. & Piggins, H. D. Vasoactive intestinal polypeptide (VIP) phase-shifts the rat suprachiasmatic nucleus clock in vitro. Eur J Neurosci 13, 839–843 (2001). 10.1046/j.0953-816x.2000.01437.x

54 Watanabe, K., Vanecek, J. & Yamaoka, S. In vitro entrainment of the circadian rhythm of vasopressin-releasing cells in suprachiasmatic nucleus by vasoactive intestinal polypeptide. Brain Res 877, 361–366 (2000). 10.1016/s0006-8993(00)02724-4

55 Piggins, H. D., Antle, M. C. & Rusak, B. Neuropeptides phase shift the mammalian circadian pacemaker. J Neurosci 15, 5612–5622 (1995). 10.1523/JNEUROSCI.15-08-05612.1995

56 Xie, L. et al. Cholecystokinin neurons in mouse suprachiasmatic nucleus regulate the robustness of circadian clock. Neuron 111, 2201–2217 e2204 (2023). 10.1016/j.neuron.2023.04.016

57 Dana, H. et al. High-performance calcium sensors for imaging activity in neuronal populations and microcompartments. Nat Methods 16, 649–657 (2019). 10.1038/s41592-019-0435-6

58 Chien, Y. F. et al. Dual GRIN lens two-photon endoscopy for high-speed volumetric and deep brain imaging. Biomed Opt Express 12, 162–172 (2021). 10.1364/BOE.405738

59 Dimidschstein, J. et al. A viral strategy for targeting and manipulating interneurons across vertebrate species. Nat Neurosci 19, 1743–1749 (2016). 10.1038/nn.4430

60 DeNardo, L. A. et al. Temporal evolution of cortical ensembles promoting remote memory retrieval. Nat Neurosci 22, 460–469 (2019). 10.1038/s41593-018-0318-7

61 Guenthner, C. J., Miyamichi, K., Yang, H. H., Heller, H. C. & Luo, L. Permanent genetic access to transiently active neurons via TRAP: targeted recombination in active populations. Neuron 78, 773–784 (2013). 10.1016/j.neuron.2013.03.025

62 Ji, N., Freeman, J. & Smith, S. L. Technologies for imaging neural activity in large volumes. Nat Neurosci 19, 1154–1164 (2016). 10.1038/nn.4358

63 Mitani, A. & Komiyama, T. Real-Time Processing of Two-Photon Calcium Imaging Data Including Lateral Motion Artifact Correction. Front Neuroinform 12, 98 (2018). 10.3389/fninf.2018.00098

64 Stringer, C., Wang, T., Michaelos, M. & Pachitariu, M. Cellpose: a generalist algorithm for cellular segmentation. Nat Methods 18, 100–106 (2021). 10.1038/s41592-020-01018-x

65 Pachitariu, M. & Stringer, C. Cellpose 2.0: how to train your own model. Nat Methods 19, 1634–1641 (2022). 10.1038/s41592-022-01663-4

66 Grapengiesser, E. Cell photodamage, a potential hazard when measuring cytoplasmic Ca2+ with fura-2. Cell structure and function 18, 13–17 (1993).

67 Knight, M. M., Roberts, S. R., Lee, D. A. & Bader, D. L. Live cell imaging using confocal microscopy induces intracellular calcium transients and cell death. American Journal of Physiology-Cell Physiology 284, C1083–C1089 (2003).

68 Grapengiesser, E. Cell photodamage, a potential hazard when measuring cytoplasmic Ca2+ with fura-2. Cell Struct Funct 18, 13–17 (1993). 10.1247/csf.18.13

69 Knight, M. M., Roberts, S. R., Lee, D. A. & Bader, D. L. Live cell imaging using confocal microscopy induces intracellular calcium transients and cell death. Am J Physiol Cell Physiol 284, C1083–1089 (2003). 10.1152/ajpcell.00276.2002

70 Banerjee, P. K. & Toga, A. W. Image alignment by integrated rotational and translational transformation matrix. Phys Med Biol 39, 1969–1988 (1994). 10.1088/0031-9155/39/11/011

71 Stowie, A. et al. Arginine-vasopressin-expressing neurons in the murine suprachiasmatic nucleus exhibit a circadian rhythm in network coherence in vivo. Proc Natl Acad Sci U S A 120, e2209329120 (2023). 10.1073/pnas.2209329120

72 Li, J. Y. & Schmidt, T. M. Divergent projection patterns of M1 ipRGC subtypes. J Comp Neurol 526, 2010–2018 (2018). 10.1002/cne.24469

73 Albus, H., Vansteensel, M. J., Michel, S., Block, G. D. & Meijer, J. H. A GABAergic mechanism is necessary for coupling dissociable ventral and dorsal regional oscillators within the circadian clock. Curr Biol 15, 886–893 (2005). 10.1016/j.cub.2005.03.051

74 Choi, H. J. et al. Excitatory actions of GABA in the suprachiasmatic nucleus. J Neurosci 28, 5450–5459 (2008). 10.1523/JNEUROSCI.5750-07.2008

75 Churchland, M. M. & Shenoy, K. V. Preparatory activity and the expansive null-space. Nat Rev Neurosci (2024). 10.1038/s41583-024-00796-z

76 Sonoda, T. et al. A noncanonical inhibitory circuit dampens behavioral sensitivity to light. Science 368, 527–531 (2020). 10.1126/science.aay3152

