## supplemental figures and legends for "Discrete photoentrainment of mammalian central clock is regulated by bi-stable dynamic network in the suprachiasmatic nucleus"

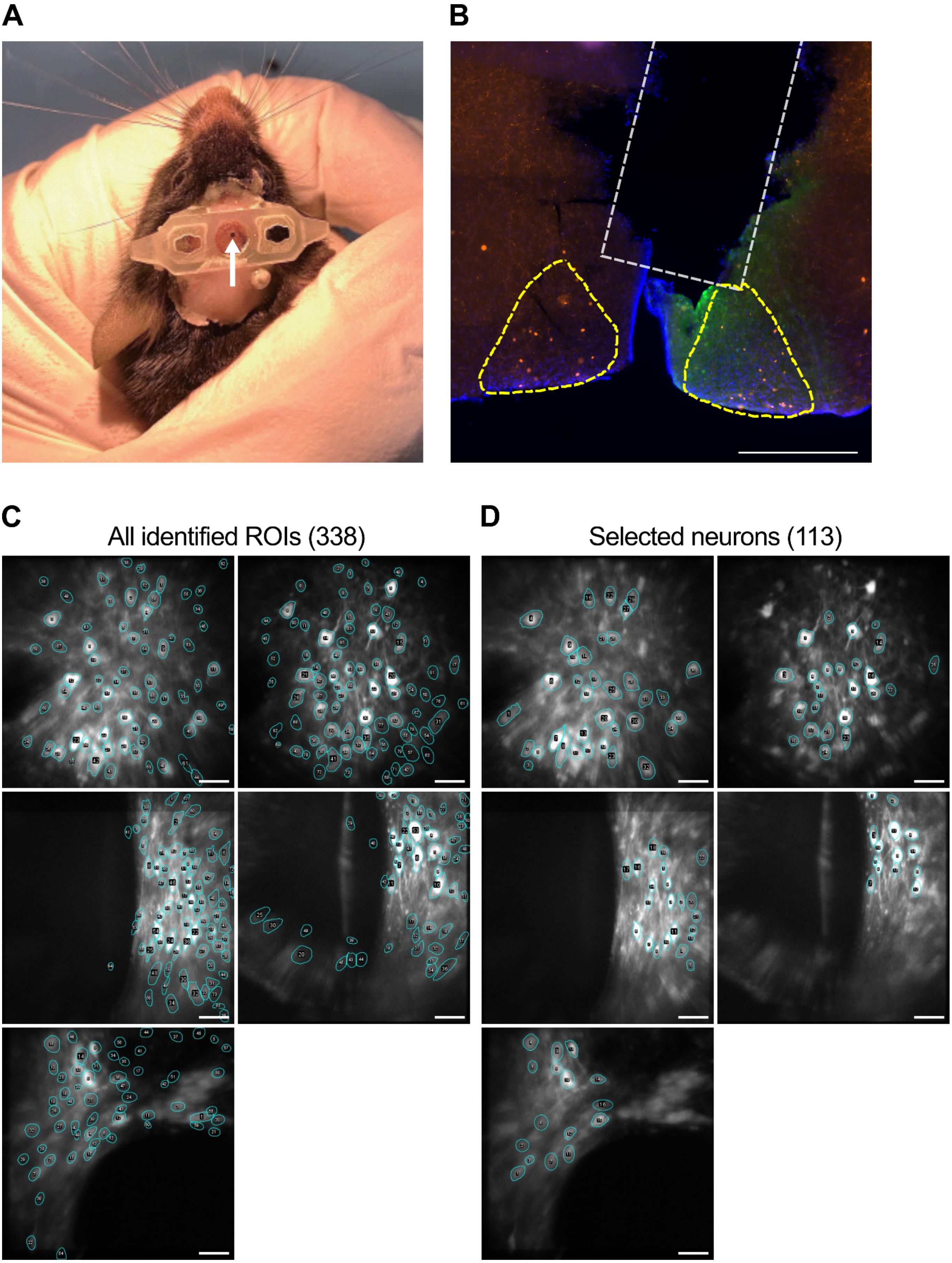
A
B
C
All identified ROIs (338)
D
Selected neurons (113)

**Extended Figure 1.**

**A.** Photo of a mouse after GRIN endoscope implantation and head bar attachment. The head bars are to protect endoscopes from damage and to connect mice and the microscope. Arrowhead indicates GRIN endoscope. **B.** Immunofluorescence staining of the SCN (indicated by yellow outline) coronal section with the trajectory of an endoscope (indicated by white outline). Orange: tdTomato, green: anti-GFP, blue: DAPI. **C.** Representative images for all 5 recording focus planes, outlines indicate ROI identified at least once in 27 trails (341 in total). **D.** Representative images for all 5 focus planes, outlines indicate neurons identified in every trial (113 in total). Scale bar: 500 µm in (B) and 100 µm in (C) and (D).

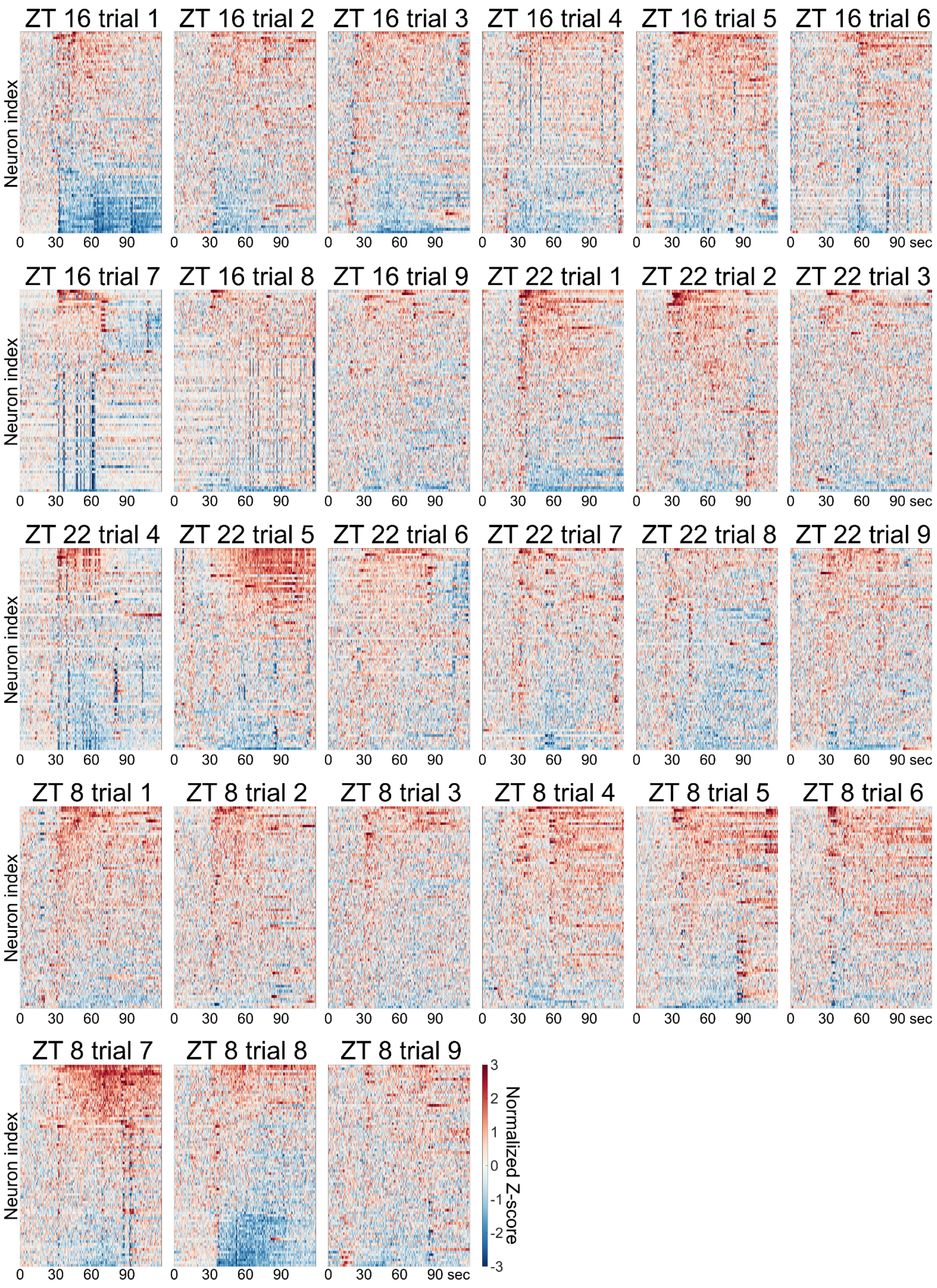


**Extended Figure 2.**

Heat map of normalized Z-score traces from 113 identified neurons in all 27 trials. Each trail is sorted with mean Z-score independently.

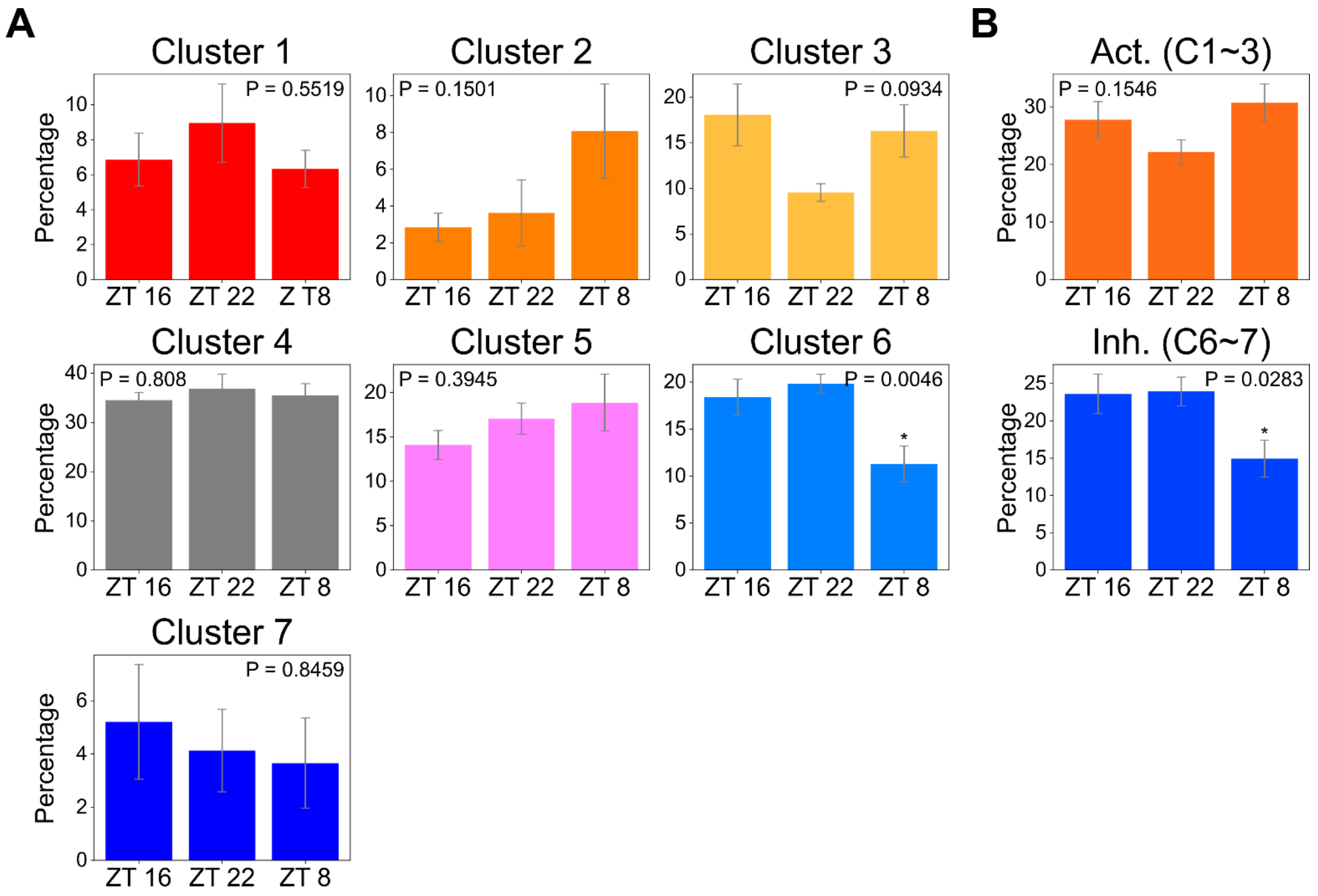


**Extended Figure 3.**

**A.** Comparing the percentage of neurons sorted into each cluster between different ZTs. **B.** Comparing the summation of activation or inhibition clusters between different ZTs. * indicates $p < 0.05$ with one-way ANOVA and Tukey post hoc tests.

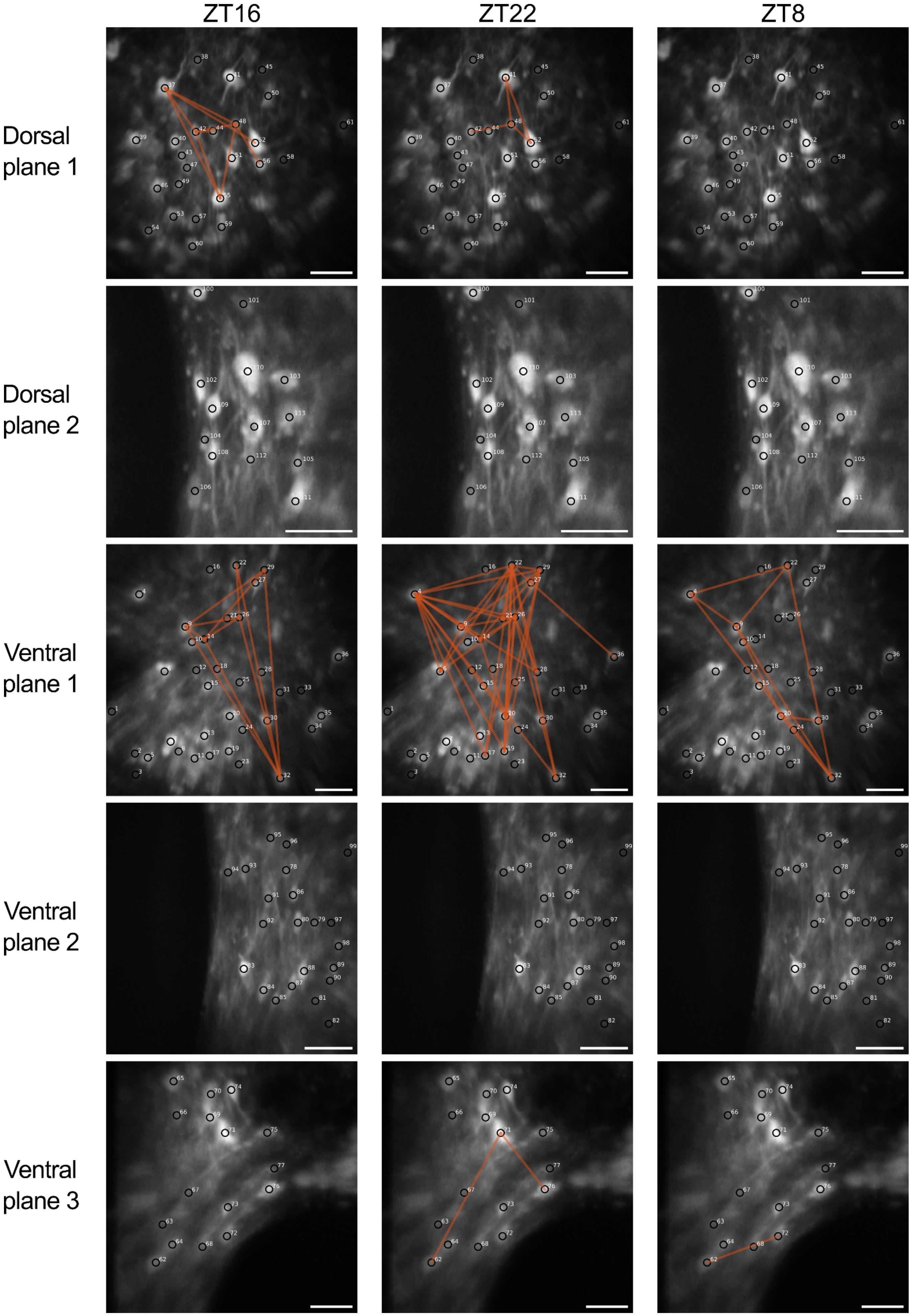
ZT16
ZT22
ZT8
Dorsal plane 1
Dorsal plane 2
Ventral plane 1
Ventral plane 2
Ventral plane 3

**Extended Figure 4.**

Highly correlated neuron pairs (average r from 9 repeats > 0.5) in all five layers from three time points are linked with orange lines. Scale bar: 100 μm.

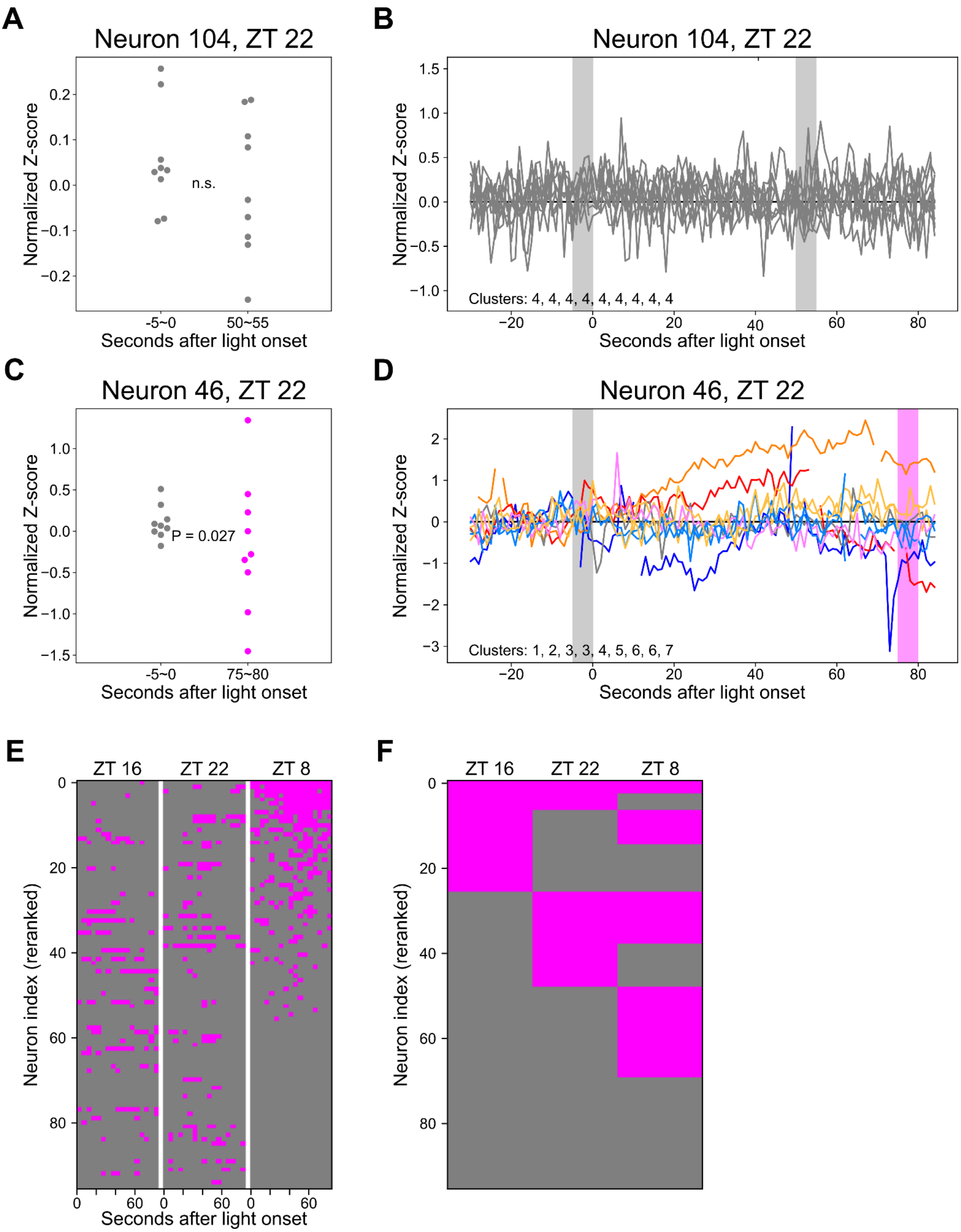
A
Neuron 104, ZT 22
Normalized Z-score
n.s.
-5~0
50~55
Seconds after light onset
B
Neuron 104, ZT 22
Normalized Z-score
Clusters: 4, 4, 4, 4, 4, 4, 4, 4, 4
Seconds after light onset
C
Neuron 46, ZT 22
Normalized Z-score
P = 0.027
-5~0
75~80
Seconds after light onset
D
Neuron 46, ZT 22
Normalized Z-score
Clusters: 1, 2, 3, 3, 4, 5, 6, 6, 7
Seconds after light onset
E
ZT 16
ZT 22
ZT 8
Neuron index (reranked)
Seconds after light onset
F
ZT 16
ZT 22
ZT 8
Neuron index (reranked)

**Extended Figure 5.**

**A & C.** Representative neurons' normalized Z-scores from nine trials showing the baseline GCaMP response (5-second average) before light onset compared with the GCaMP response 50 and 75 seconds after light onset. **B & D.** The full traces of nine trials for representative neurons. Each color represents a different light response cluster. A & B depict a representative group 3 neuron whose post-onset variances are not significantly greater than the pre-onset baseline. C & D illustrate another representative group 3 neuron with certain 5-second bins post-onset exhibiting significantly increased variance compared to the pre-onset baseline. **E.** Composite plot from group 3 neurons where significantly increased post-onset variance are highlighted in magenta. Levene's test for equality of variances, $p < 0.05$ **F.** Group 3 neurons are marked in magenta if at least one bin demonstrates significant variance in the Levene's test. This analysis indicates that 69 of the group 3 neurons (72.6%) display high variation amount trails at same ZTs.

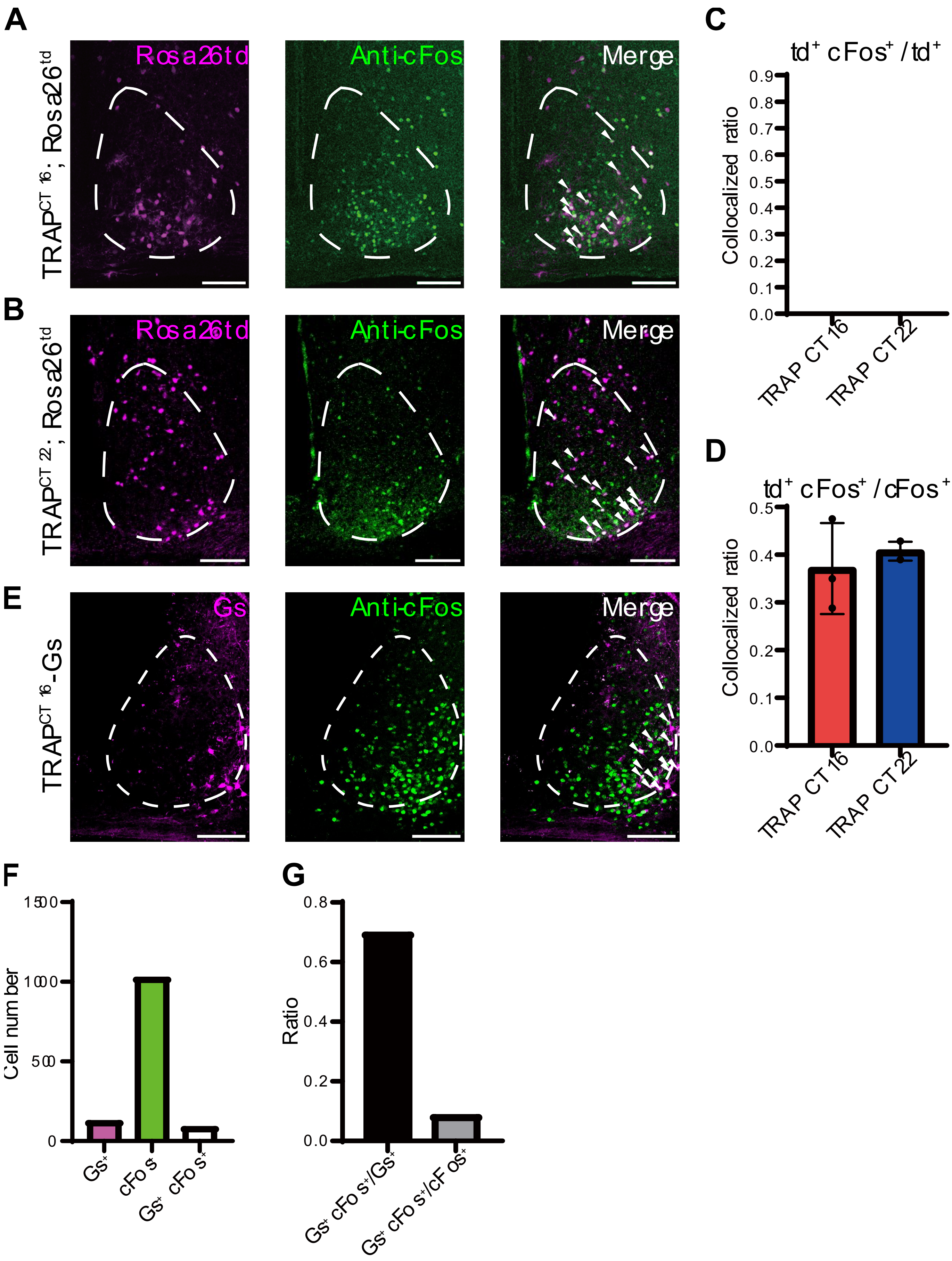

A
B
C
D
E
F
G
Rosa26td
Anti-cFos
Merge
Gs
td⁺ cFos⁺ / td⁺
td⁺ cFos⁺ / cFos⁺
Collocalized ratio
TRAP CT 16
TRAP CT 22
Cell number
Ratio
Gs⁺
cFos
Gs⁺ cFos⁺
Gs⁺ cFos⁺/Gs⁺
Gs⁺ cFos⁺/cFos⁺

**Extended Figure 6.**

**A.** Representative fluorescence images for CT 16-trapped light responsive neurons (magenta) in TRAP2;Ai14 (Rosa26td) mice and cFos immune-positive SCN neurons (green) for CT 16 light pulse. **B.** Representative fluorescence images for CT 22-trapped light responsive neurons (magenta) in TRAP2;Ai14 (Rosa26td) mice and cFos immune-positive SCN neurons (green) for CT 22 light pulse. **C.** Statistic of colocalized ratio for trapped and anti-cFos double-positive SCN neurons over trapped SCN neurons, which implies the specificity of trapped neurons. **D.** Statistic of colocalized ratio for trapped and anti-cFos double-positive SCN neurons over anti-cFos SCN neurons, which implies the efficiency of trapped neurons. (Two-way ANOVA: * $p < 0.05$, ** $p < 0.01$, *** $p < 0.001$, n=3) **E.** Representative image for DREADDs (rM3Ds)-expressing CT 16-trapped light response and cFos immune-positive SCN neurons for CT16 light pulse. **F.** Bar chart shows cell number for Gs-expressing, anti-cFos, and double positive SCN neurons in TRAP-CT 16 mice. **G.** The Bar chart shows the specificity and efficiency reflected by the colocalized ratio in the CT 16-trapped SCN neurons. Scale bar: 100 μm.

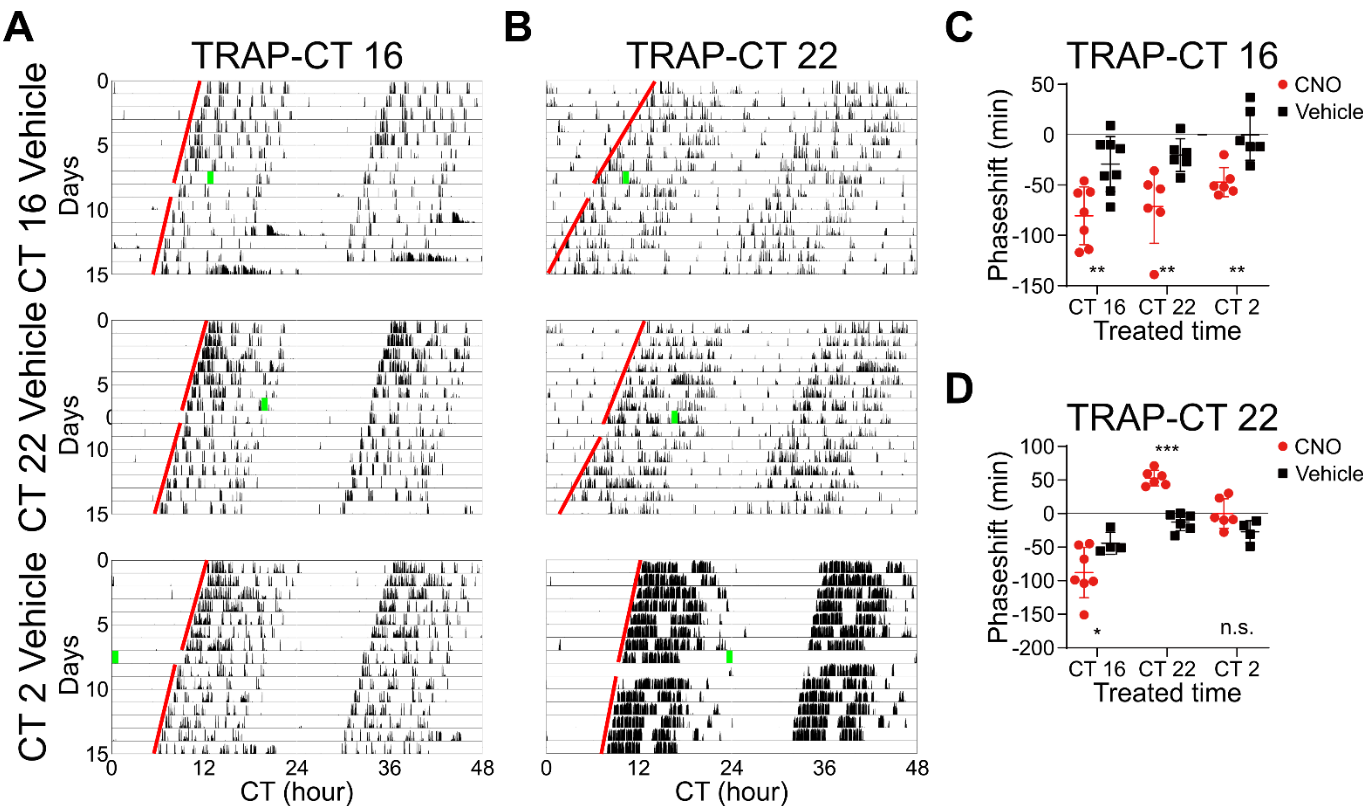


**Extended Figure 7.**

**A.** Representative actogram for DREADDs (rM3Ds)-expressing TRAP-CT 16 mice. **B.** Representative actogram for DREADDs (rM3Ds)-expressing TRAP-CT 22 mice. Green bars indicate the time points of CNO or vehicle (saline) injection, while red lines indicate linear regression line for activity onsets. **C.** Statistics of phase shift analysis for CNO and vehicle (saline) treatment in the TRAP-CT 16 mice. Two-way ANOVA, * p < 0.05, ** p < 0.01, *** p < 0.001, vehicle group, n= 6-8; CNO group, n=6-7. **D.** Statistics of phase shift analysis for CNO and vehicle treatment in the TRAP-CT 22 mice. Two-way ANOVA: * p < 0.05, ** p < 0.01, *** p < 0.001, vehicle group, n=4-6; CNO group, n=6-7.
